# A pyoverdine-type metallophore is required for lanthanide-dependent growth in *Pseudomonas putida*

**DOI:** 10.64898/2026.09.08.750064

**Authors:** Kalen Rasmussen, Jonathan Michael, Alex F. Benson, Kelsey J. Ramirez, Morgan A. Ingraham, Allegra T. Aron, Allison Werner

## Abstract

Lanthanides (Ln) are recently discovered cofactors for alcohol metabolism in a growing number of bacteria, including the soil-dwelling bacterium *Pseudomonas putida,* yet the mechanisms for Ln uptake are poorly defined. Here, we discovered that the non-ribosomal peptide synthase biosynthetic gene cluster, *pvdLIJD*, was essential to Ln-dependent growth in *P. putida*. Pyoverdine G4R and its precursor ferribactin were identified as *pvdLIJD* gene products with La-responsive abundances, and both chelated La as well as iron. Deletion of the putative pyoverdine outer membrane transport system encoded by *fpvA* and *exbBD:tonB* were strongly linked to Ln-dependent growth in a concentration-dependent manner. Global transcriptomes were remodeled in response to La, but weak evidence of transcriptional *pvdLIJD* regulation and no evidence of discernment between La provided as chloride versus oxide were observed. Overall, this work establishes pyoverdines as metallophores involved in La uptake, re-framing them from strictly iron-scavenging siderophores to dual-purpose metallophores with Ln-binding activity.

## INTRODUCTION

Metal ions are required for a diverse range of biological functions (*1*, *2*), with transition metals such as copper, cobalt, manganese, and iron (Fe) being well-established metalloprotein cofactors (*3–5*). While metalloprotein metal recognition can be highly-specific (*6*), the stability of copper and, to a lesser extent, zinc complexes is generally stronger than other transition metals (*7*). Thus, microorganisms must ensure proper metalation and maintain biological function through homeostatic processes for spatiotemporal control of metal ions, including uptake, trafficking, and storage (*8*).

Microorganisms secrete chemically-diverse metal-chelating metabolites (“metallophores”) to facilitate uptake and increase the bioavailability of extracellular metals (*9–11*). High-affinity Fe-binding (siderophore) systems are thought to overcome the low solubility of Fe in natural environments (*12*), though siderophores have an increasingly recognized importance in metal-rich environments (*13*, *14*). Metallophore biosynthetic gene clusters (BGCs) encode functions including biosynthesis, export, import of metal-metallophore complexes, intracellular metal release, and BGC regulation (*15*). Gram-negative bacteria often utilize substrate-specific outer membrane receptors (OMR) coupled to the inner membrane TonB-ExbB-ExbD energy-transducing system for import (*16*, *17*), metal reduction or protein-mediated ligand exchange for periplasmic release (*18*), and ATP-binding cassette (ABC)-type transport into the cytoplasm (*19*, *20*).

Lanthanides (Lns), a series of 15 consecutive *f*-block elements with abundances up-to 100 mg/kg in soil (*21*), have recently emerged as biologically relevant metals. In 2011, Hibi *et al.* demonstrated the phyllospheric bacterium *Methylobacterium radiotolerans* utilized a lanthanum (La) cofactor for methanol metabolism via the pyrroloquinoline quinone (PQQ)-dependent methanol dehydrogenase XoxF (*22*). Studies since have sought to understand Ln homeostasis in *Methylobacteriacae* and have revealed selective Ln-binding proteins (*23–25*), including those with potential biotechnological applications for critical mineral recovery (*26–28*). The full mechanisms of Ln homeostasis are still largely unresolved, but the use of Ln-chelating metallophores (“lanthanophores”) for scavenging has been proposed (*29*, *30*).

Citrate-based polycarboxylate lanthanophores have been reported in *Methylobacterium* (*31*, *32*), and deletion of the BGC encoding the polycarboxylate siderophore staphyloferrin B in the facultative methylotroph *M. aquaticum* 22A stunted but did not altogether prevent Ln-dependent growth (*31*). Methylolanthanin, a polycarboxylate metallophore with two terminal 4-hydroxybenzoate groups, was discovered in *Methylobacterium extorquens* AM1 and found to bind Lns by metal infusion mass spectrometry (MS) (*32*). Deletion of the methylolanthnin BGC, *mll,* stunted growth under the tested conditions, and *mll* was upregulated during cultivation on Nd_2_O_3_ as compared to NdCl_3_, leading the authors to propose methylolanthanin mediates Ln solubilization, uptake in association with the OMR MluA, and coordination with XoxF (*32*).

*Pseudomonas putida* KT2440 (hereafter *P. putida*) is one of the few non-methylotrophic bacteria reported to utilize Ln-dependent metabolism. Wehrmann *et al.* established this in a series of seminal reports, first showing *P. putida* utilizes the Ln-dependent PQQ alcohol dehydrogenase (ADH) PedH for metabolism of aromatic and aliphatic alcohols (**Fig. 1A**) (*33*). PedH is proposed to be largely redundant with the calcium (Ca)-dependent PQQ ADH PedE (*33–35*), and expression of *pedE* and *pedH* are modulated through a partially-resolved Ln-responsive regulatory network termed the “REE-switch” (*36*). Deletion of the inner membrane ABC-type transporter PedA_1_A_2_BC resulted in a strain that required saturating LaCl_3_ (≥10 µM), suggesting cytoplasmic La uptake is involved in Ln-dependent growth (*37*). While many knowledge gaps in *P. putida* Ln homeostasis remain unresolved, the wealth of accumulated knowledge and genetic engineering tools support progress, with strong relevance to *P. putida*’s widespread application in biomass and plastics valorization (*38–41*).

**Figure 1.**
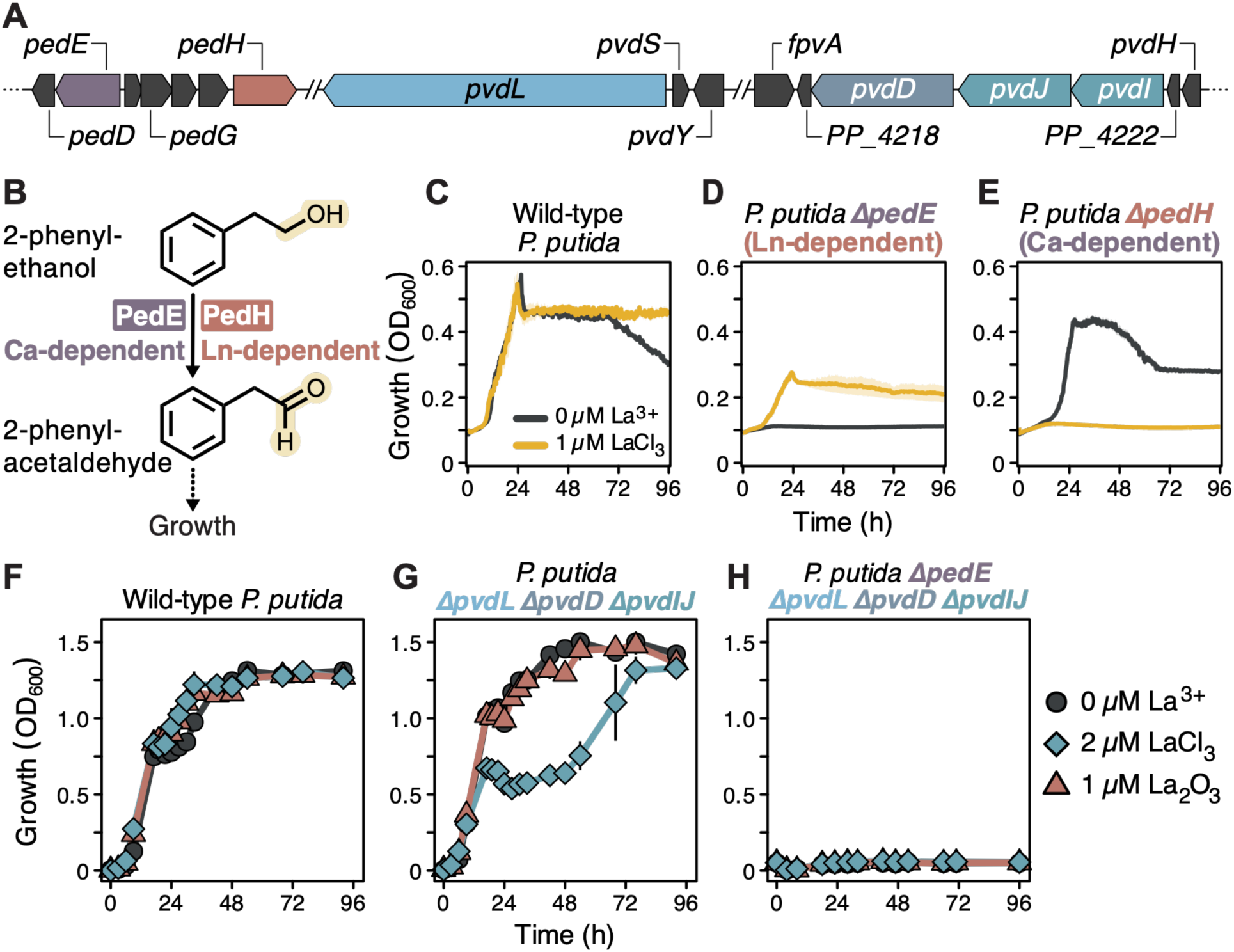
Genetic organization and growth profiles of *P. putida pedE, pedH,* and *pvdLIJD* deletion mutants. (**A**) Schematic of the genetic organization and flanking regions of *pedE*, *pedH*, and associated NRPS genes within the *ped* and *pvd* biosynthetic gene clusters. (**B**) *P. putida* harbors two PQQ-dependent ADHs, the La-dependent PedH and Ca-dependent PedE, which mediate conversion of 2-phenylethanol (2-PE) to 2-phenylaldehyde. (**C**) Wild-type *P. putida*, (**D**) *P. putida* Δ*pedE*, or (**E**) *P. putida* Δ*pedH* growth during cultivations in modified MP media supplemented with 5 mM 2-PE in 200 µL honeycomb microtiter plates at 30°C. Media was supplemented with LaCl_3_ as indicated and data represent the mean values of biological triplicates; shaded regions correspond to standard deviation. (**F**) Wild-type *P. putida*, (**G**) *P. putida* Δ*pvdL* Δ*pvdD* Δ*pvdIJ,* or (**H**) *P. putida* Δ*pedE* Δ*pvdL* Δ*pvdD* Δ*pvdIJ* growth during cultivations in 60 mL modified MP medium in 250 mL acid-washed polycarbonate flasks at 30°C and 225 rpm. Data points represent the mean values of biological triplicates and error bars correspond to standard deviations.

Here, we report that pyoverdine-type metallophores are required for lanthanide (Ln)-dependent growth in *P. putida*. Deletion of the pyoverdine-encoding BGC *pvdLIJD* had negligible effects in wild-type *P. putida*, but abolished growth in a *pedE* deletion strain. We identified the *pvdLIJD*-encoded metabolites as pyoverdine G4R and its precursor ferribactin, and directly observed La-chelation by both compounds using metal infusion MS. The global transcriptome and *pvdLIJD* expression were sensitive to La supplementation, but did not differentiate between chloride and oxide forms. Gene deletions of the OMR FpvA and the TonB-ExbBD energy transduction system in a Ln-dependent background strain were detrimental in a *pedE* deletion background in a concentration-dependent manner. In sum, this work demonstrates that *P. putida* utilizes pyoverdine G4R and/or ferribactin for La chelation and uptake, establishing a new class of bacterial metallophores that enable La-dependent carbon metabolism.

## RESULTS

### Defined growth conditions isolate Ln-dependent PedH metabolism of 2-phenylethanol

We first sought to establish the genetic backgrounds under which La uptake is essential for growth. *P. putida* encodes both a Ca-dependent PQQ ADH, *pedE*, and Ln-dependent PQQ ADH, *pedH* (**Fig. 1A**). 2-Phenylethanol (2-PE) was selected as the carbon and energy source as *P. putida* has been previously shown to grow via the PQQ ADH pathways (**Fig. 1B**). (*33*). We deleted *pedE* to constrain metabolism to the PedH- and Ln-dependent pathway (*P. puitda* Δ*pedE*), or deleted *pedH* to constrain metabolism to the PedE- and Ca-dependent pathway (*P. putida* Δ*pedH*) (**Table S1**). A double ADH mutant (*P. putida* Δ*pedE* Δ*pedH*) was also constructed.

To determine the optimal media composition for isolating Ln-dependent growth, we evaluated growth of wild-type *P. putida* and the ADH deletion mutants with varied buffers and trace metal concentrations. Standard M9 minimal media contains high phosphate concentrations, where La^3+^ readily precipitates (*42*, *43*). To avoid this, modified M9 and MP media use elevated concentrations of Ca in conjunction with trace metals supplementation and Na_3_-citrate to promote metal solubility (**Table S2**), (*32*, *33*, *36*) . Additionally, MP media uses PIPES buffer and is commonly supplemented with a “C7” metals mixture containing micromolar concentrations of Co, Cu, Fe, Mn, Mo, Ni, W, and Zn (*33*, *37*) (**Table S3**). While PIPES didn’t support growth as a sole carbon source, the expected phenotype of ADH mutants was disrupted, whereas a a 8.5 mM phosphate buffer in MP media maintained pH ∼7 over 113 h (**Figs. S1-2**).

The ADH mutants were cultivated on 5 mM 2-PE in Modified M9 versus MP media with varying LaCl_3,_ CaCl_2_, C7 metals, and FeSO_4_ concentrations (**Fig. S3-6**). *P. putida* Δ*pedE* Δ*pedH* did not grow regardless of media or La^3+^ concentration (**Fig. S3**), confirming no other ADHs in *P. putida* are capable of 2-PE metabolism. As expected, wild-type *P. putida* required either Ca or La for growth (**Fig. 1C**) and exhibited impaired growth in the absence of C7 metals, with minimal differences in Modified M9 versus MP medias (**Figs. S4-5**). As expected, *P. putida* Δ*pedE* required La for growth (**Fig. 1D**), regardless of La concentration, base media, or C7 supplementation (**Figs. S4-5**). *P. putida* Δ*pedH* grew in the absence of La and no growth was observed with 1 μM LaCl_3_ (**Fig. 1E**), consistent with metabolism via the Ca-dependent PedE and the influence of the REE-switch on expression of *pedE* (*33*, *36*). Both *P. putida* Δ*pedE* and *P. putida* Δ*pedH* exhibited an extended lag phase at 0.1 but not 10 μM LaCl_3_ in the presence of C7 metals (**Fig. S5**), suggesting a dose-dependent effect of transition metals on the REE-switch, as has been shown for high concentrations of Fe (*37*). Removing all trace metals except Fe minimized the regulatory effects imparted by the C7 metals mixture while still maintaining growth, with 1 µM FeSO_4_ being optimal (**Fig. S6**). Given these results, we proceeded using 5 mM 2-PE, 8.5 mM phosphate buffer (pH 7), 0.5 mM MgCl_2_, 100 µM CaCl_2_, 8 mM (NH_4_)_2_SO_4_, 45 µM Na_3_-citrate, and 1 µM FeSO₄-7H₂O, which we hereafter refer to as “modified MP medium”.

### The non-ribosomal peptide synthase (NRPS) biosynthetic gene cluster (BGC) *pvdLIJD* is required for La-dependent growth on 2-PE

Having established conditions under which Ln uptake is essential for growth, we next sought to investigate the possibility of a La-chelating secondary metabolite in *P. putida*. The bioinformatic tool antiSMASH v8 (*44*), which facilitates genome-wide identification of genes encoding metallophore systems, was used to mine the *P. putida* genome for putative secondary metabolites and their corresponding BGCs. Ten putative regions were identified (**Table S4**), including a non-ribosomal peptide synthase (NRPS) region associated with a pyoverdine-type metabolite associated with Fe acquisition in pseudomonads (*45*, *46*).

Based on homology to *P. aeruginosa,* four pyoverdine-associated NRPS genes – *pvdL*, *pvdI*, *pvdJ*, and *pvdD* – are present in *P. putida*’s genome and are intertwined with the *fpv* gene cluster for Fe scavenging (**Fig. 1A**). In *P. aeruginosa*, NRPS proteins sequentially synthesize pyoverdine in order of PvdL, PvdI, PvdJ, and PvdD (11). Previously, the effect of *pvdD* deletion in *P. putida* was shown to be inconsequential to Ln-dependent growth (*37*). However, if biosynthesis follows that of *P. aeruginosa, P. putida* Δ*pvdD* would produce a truncated pyoverdine with unknown chelation attributes.

Given these results, we hypothesized that *pvdL*, *pvdI*, *pvdJ*, and *pvdD* together produced a Ln-chelating metabolite necessary for Ln-dependent growth. We constructed a four-gene deletion mutant (*P. putida* Δ*pvdL* Δ*pvdD* Δ*pvdIJ*) in wild-type and Δ*pedE* backgrounds (**Table S1**) and cultivated all three strains on 5 mM 2-PE in the absence of La^3+^, 1 µM La_2_O_3_, or 2 µM LaCl_3_ (**Fig. S8**). Wild-type *P. putida* grew well irrespective of La supplementation (**Fig. 1F**). *P. putida* Δ*pvdL* Δ*pvdD* Δ*pvdIJ* grew under all conditions, but growth on 2 µM LaCl_3_ displayed biphasic profile, lagging ∼36 h after an initial period of growth (**Fig. 1G**). Stacking in Δ*pedE*, necessitating Ln uptake into the periplasm for PedH-dependent growth, with Δ*pvdL* Δ*pvdD* Δ*pvdIJ* resulted in complete abolishment of growth with either LaCl_3_ or La_2_O_3_ supplementation (**Fig. 1H**). These results demonstrate the requirement of the *pvdLIJD* BGC for PedH-dependent growth on 2-PE in *P. putida*.

### The *pvdLIJD*-encoded metallophores pyoverdine G4R and its ferribactin precursor chelate Fe and La

We next sought to confirm the *pvdLIJD-*encoded metabolite was a pyoverdine, and determine whether its abundance was affected by the presence and form of supplemented La. We cultivated wild-type *P. putida*, *P. putida* Δ*pedE*, and *P. putida* Δ*pvdL* Δ*pvdD* Δ*pvdIJ* in modified MP media supplemented with 5 mM 2-PE in the absence of La, with 1 µM La_2_O_3_, or with 2 µM LaCl_3_, and analyzed the supernatant for the presence of pyoverdines previously reported in *P. putida* (*47–49*).

We detected the exact mass of a pyoverdine previously reported by El Din *et al.* (*47*) and Baune *et al*. (*48*) (1073.3289 m/z), referred to as G4R and G4R-A, respectively. The pyoverdines reported as G4R-A and G4R have a succinamide side-chain and are otherwise chemically identical; we henceforth refer to this pyoverdine as G4R. Examination of the fragmentation pattern of the detected adducts agreed with previously described b-, c-, and y-ions, confirming the identity as G4R (**Figs. 2A & S9A, Table S5**) (*48–50*). The cyclic peptide variant previously reported during growth on succinate (*49*) was not detected.

**Figure 2.**
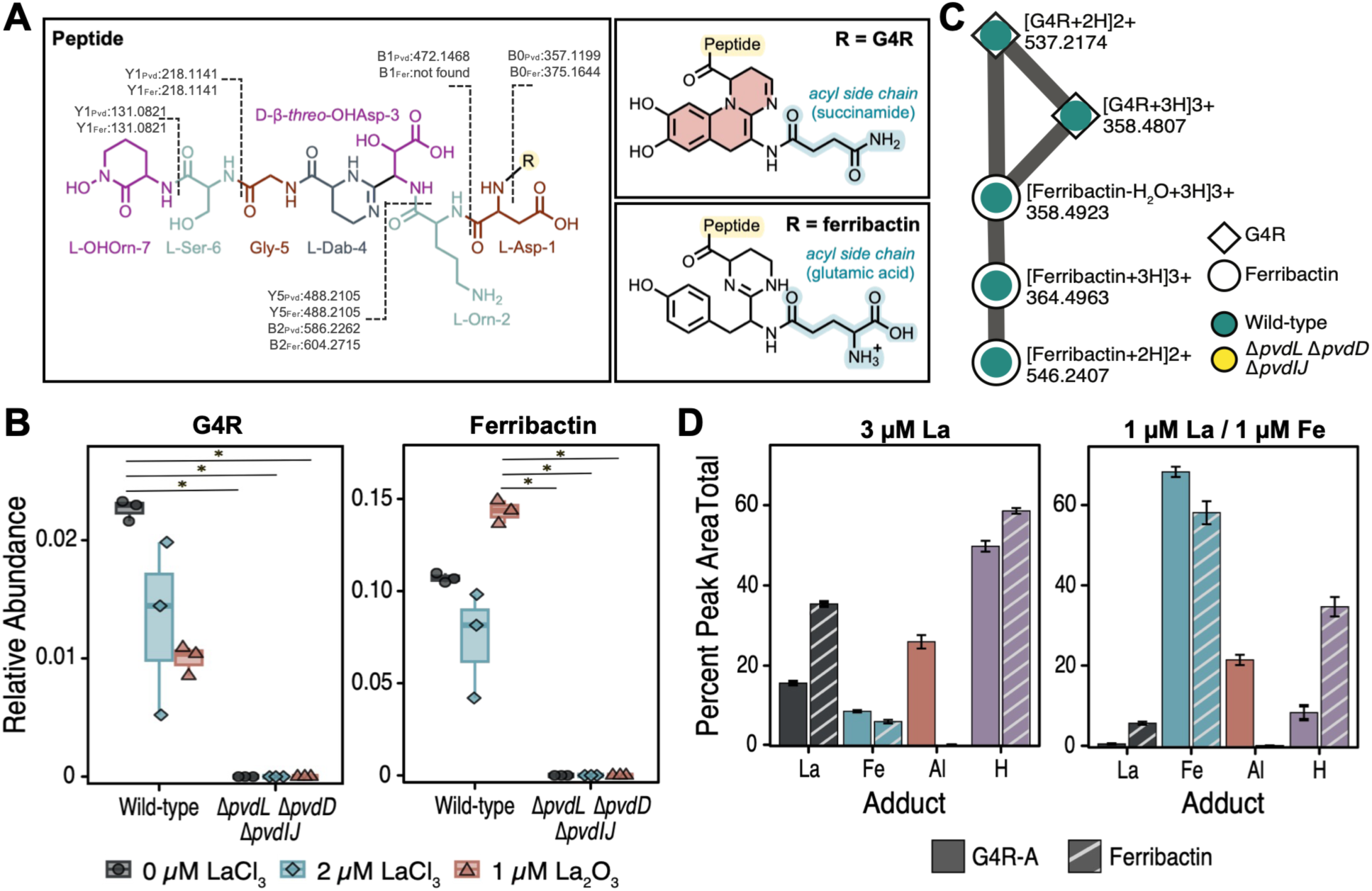
Relative abundance and metal chelation of pyoverdine G4R and ferribactin in wild-type *P. putida* and *P. putida* Δ*pvdL* Δ*pvdIJ* Δ*pvdD*. (**A**) Proposed structures of pyoverdine G4R, as reported previously (*47*, *48*), and ferribactin. (**B**) Relative abundance of pyoverdine G4R and ferribactin molecules in the supernatants of wild-type *P. putida* and *P. putida* Δ*pvdL* Δ*pvdIJ* Δ*pvdD* cultivations in modified MP media supplemented with 5 mM 2-PE and La as indicated. Relative abundance based on blank-subtracted TIC; significance calculated by Kruskal-Wallis with Dunn’s test and BH correction (*, *p*<0.05). (**C**) Feature-based molecular network of the detected pyoverdine G4R and ferribactin molecules. Pie charts represent the proportion of peak areas found in each sample type; symbol shapes indicate the molecule. (**D**) Percent of peak areas by adduct type measured after infusion with 3 µM LaCl_3_ or a mixture of 1 µM LaCl_3_ and 1 µM FeCl_3_. Adducts are singly, doubly, and triply charged variants of each type. Error bars represent one standard error from the mean of 2 technical replicates of 3 biological replicates (La) or one standard error from the mean of 3 technical replicates of 3 biological replicates (La/Fe).

Pyoverdine G4R and its addunts were absent in *P. putida* Δ*pvdL* Δ*pvdIJ* Δ*pvdD* cultivations (**Fig. 2B, S10**), confirming PvdLIJD-dependent biosynthesis. In wild-type *P. putida,* the highest relative abundance was observed in La-free media, with both LaCl_3_ and La_2_O_3_ supplementation resulting in decreased abundance (**Fig. 2B**), consistent with trends observed upon Fe-supplementation (*51*).

Next, we searched for metabolites associated with pyoverdine G4R using feature-based molecular networking combined with a MassQL (*52*) query tailored to the fragmentation of pyoverdine, isopyoverdine, and ferribactin precursors (*53*). We identified an additional molecule characteristic of ferribactin, a biosynthetic intermediate of mature pyoverdines, networked with pyoverdine G4R (**Fig. 2C**). The molecule had an exact mass of m/z 1091.4741 with the diagnostic ferribactin m/z 136.0757 fragment (**Figs. 2A & S9B, Table S5**). Analysis of MS2 spectra identified conserved y-ions with pyoverdine G4R, but a mass deviation of 18.0463 in the b-ions attributed to the chromophore region, leading us to propose the identity as ferribactin (**Figs. 2A & S9B, Table S5**). Like pyoverdine G4R, the molecule and its adducts were not detected in the supernatant of *P. putida* Δ*pvdL* Δ*pvdD* Δ*pvdIJ* cultivations (**Figs. 2B & S10**). Unlike pyoverdine G4R, the relative abundance of ferribactin in wild-type cultures was highest with La_2_O_3_ supplementation (**Figs. 2B & S10**).

To determine if pyoverdine G4R or ferribactin chelate La, we leveraged metal infusion MS (*54*). Wild-type samples were subjected to chromatographic separation with post-column infusion of either pure LaCl_3_ or an equimolar mixture of FeCl_3_ and LaCl_3_, and metal adducts were quantified by comparing peak areas corresponding to metal adduct masses at the same retention time. For pyoverdine G4R, La infusion resulted in 14-17% La-bound molecule, which decreased to 0.5-2% in the presence of Fe (**Fig. 2D**).

Interestingly, ferribactin complexed more readily with La: La infusion resulted in 36% La-bound molecule, decreasing to 7% La adduct peak areas in the presence of Fe, triple that of the pyoverdine G4R (**Fig. 2D**).

### The FpvA-TonB-ExbBD transport system is involved in, but not essential for, uptake of Ln-chelate complexes

We next sought to identify the transport mechanism of the Ln-pyoverdine complex, starting with antiSMASH-predicted (*44*) transport-related gene candidates. Thirty-two arrayed transposon insertion mutants (**Table S6**) (57) were grown on 2-PE in the absence or presence of 2 μM LaCl_3_, ten of which displayed changes in lag or growth rate (**Fig. S11A**). The insertion mutant in the gene encoding the OMR FpvA (*fpvA*::Tn, *PP_4217*) displayed a significant lag increase upon La addition (*p*=0.03, **Fig. S11B**). FpvA is an uncharacterized homolog of FpvA found in *P. aeruginosa*, which has been shown to work in concert with the TonB-ExbBD energy transducing system to import Fe-pyoverdine structures across the outer membrane (*16*, *56*).

To evaluate the impact of these genes on Ln-dependent growth, clean deletion mutants were constructed for *fpvA* and *exbBD:tonB* in both wild-type *P. putida* and *P. putida* Δ*pedE*. Since the FpvA-TonB-ExbBD system is involved in Fe uptake, an essential trace element, we first profiled growth on 0-50 μM FeSO_4_ to determine the Fe concentrations that permitted mutant growth but minimize interference in the presence of 2 μM LaCl_3_ (**Fig. S12**). Growth of *P. putida* Δ*pedE* Δ*fpvA* and *P. putida* Δ*pedE* Δ*exbDB:tonB* was limited but not fatal at 500 nM FeSO_4_.

Wild-type, Δ*fpvA*, Δ*tonB:exbBD*, and Δ*pedE* permutations were cultivated on 5 mM 2-PE plus 500 nM Fe at seven concentrations across 0-10 µM LaCl_3_ to examine concentration-dependent roles of FpvA, TonB, and ExbBD on La-dependent growth (**Figs. 3 & S13A**). In the absence of La, wild-type *P. putida*, *P. putida* Δ*fpvA*, and *P. putida* Δ*tonB:exbBD* displayed similar growth profiles. However, growth profiles diverged from wild-type upon La addition for both *fpvA* and *tonB:exbBD* deletion strains.

**Figure 3.**
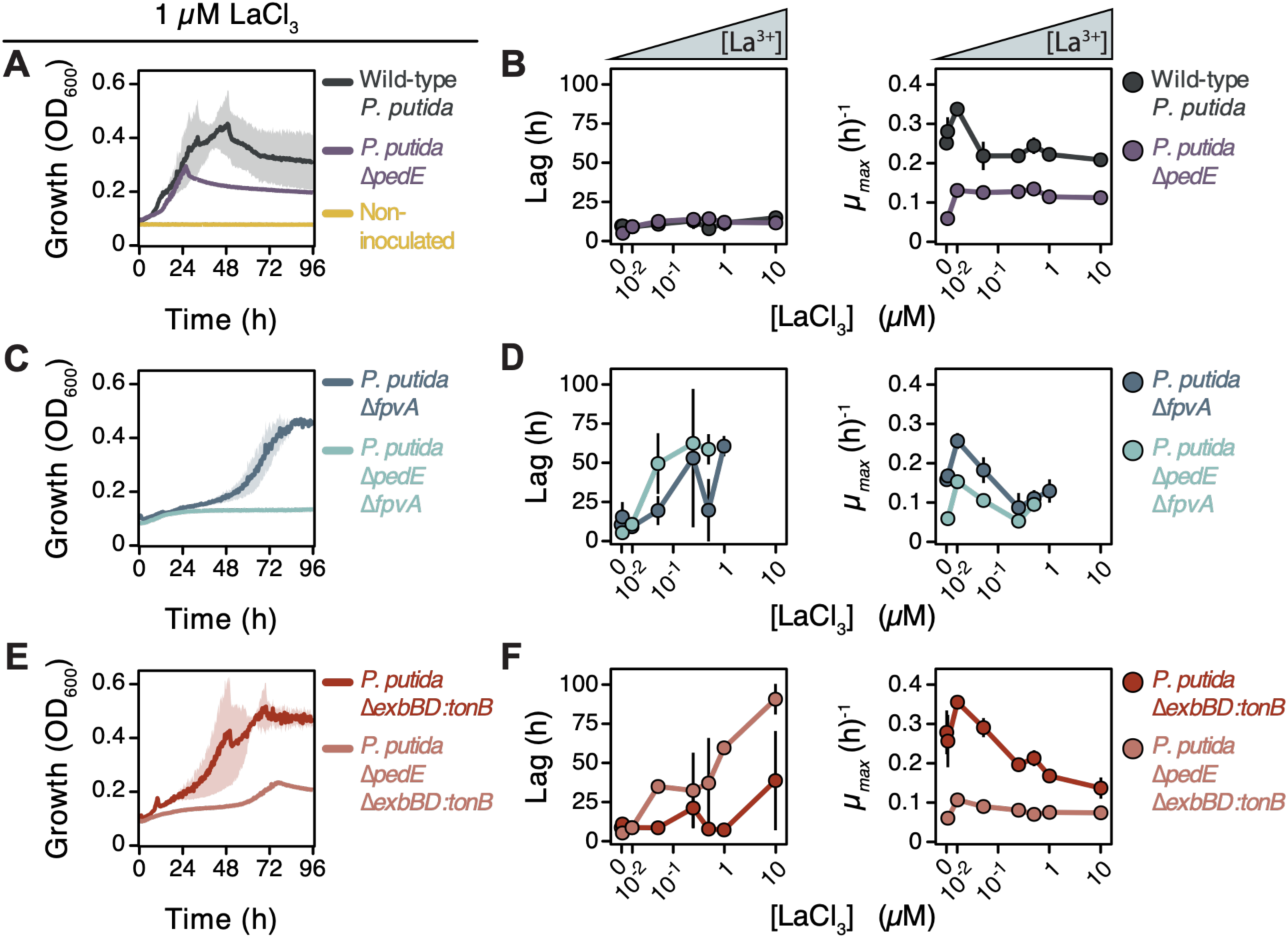
Growth profiles of *P. putida fpvA* and *exbBD:tonB* deletion mutants in wild-type and PedH-dependent backgrounds. Wild-type *P. putida* and *P. putida* Δ*pedE* control cultivations on 5 mM 2-PE supplemented with (**A**) 1 μM LaCl_3_ (growth curves) or (**B**) 0-10 μM LaCl_3_ (lag and growth rate). OMR *fpvA* deletion mutant (*P. putida* Δ*fpvA* and *P. putida* Δ*pedE* Δ*fpvA*) cultivations on 5 mM 2-PE supplemented with (**C**) 1 μM LaCl_3_ (growth curves) or (**D**) 0-10 μM LaCl_3_ (lag and growth rate). TonB-ExbBD energy transducing system deletion mutants (*P. putida* Δ*exbBD:tonB* and *P. putida* Δ*pedE* Δ*exbBD:tonB*) cultivations on 5 mM 2-PE supplemented with (**E**) 1 μM LaCl_3_ (growth curves) or (**F**) 0-10 μM LaCl_3_ (lag and growth rate). All cultivations contained 500 nM FeSO_4_. Cultures were grown in 200 µL modified MP medium at 30°C in 100-well honeycomb plates. Data points/bars represent the mean values of biological triplicates and error bars/shaded areas correspond to standard deviations. Cultures where no growth was observed were omitted from lag and growth rate calculations. See **Fig. S13** for Welch’s t-test and the Benjamini-Hochberg adjusted *p*-values for relevant comparisons.

At 1 µM LaCl₃, wild-type and *P. putida* Δ*pedE* exhibited comparable lag phases (11.46 ± 5.2 h and 11.99 ± 1.5 h, respectively) (**Figs. 3A-B & S13**). In comparison, *P. putida* Δ*fpvA* exhibited a decreased growth rate and prolonged lag phase relative to wild-type (**Figs. 3C-D & S13**). These trends were further exacerbated as the concentration of LaCl_3_ increased (**Fig. 3D**), and no growth within the 96 h study was observed at 10 µM LaCl_3_. When stacked into *P. putida* Δ*pedE*, growth was more severely impacted, with minimal impacts at nanomolar concentration but no growth observed at ≥1 µM LaCl_3_ (**Figs. 3C-D & S13**).

*P. putida* Δ*pedE* Δ*tonB:exbBD* showed reduced growth rate and increased lag at 1 µM LaCl_3_ (**Fig. 3E**), and these trends were exacerbated in a concentration-dependent manner with significant reductions in growth rate and increased in lag as La concentrations were increased, though measurable growth was still observed (**Figs. 3F & S13**). Together, these data demonstrate a clear link between FpvA-TonB-ExbBD and Ln-dependent growth, but *P. putida* Δ*pedE* Δ*fpvA* and *P. putida* Δ*pedE* Δ*tonB:exbBD* were still capable of marginal growth at concentrations ≤500 nM La, indicating the presence of a redundant mechanism for Ln outer membrane transport.

### Transcriptome expression was sensitive to La supplementation but did not distinguish between chloride and oxide forms

Given the strong decrease in pyoverdine metabolite abundance upon addition of La, we sought to understand whether the *pvd* gene cluster was transcriptionally responsive to La. Previously, it’s been indicated that LnCl_3_ rapidly dissociates into soluble Ln^3+^ whereas La from Ln_2_O_3_ remain as sparingly soluble minerals (*32*), but this is likely media-specific. We cultivated wild-type *P. putida, P. putida* Δ*pedE, P. putida* Δ*pvdL* Δ*pvdD* Δ*pvdIJ* and *P. putida* Δ*pedE* Δ*pvdL* Δ*pvdD* Δ*pvdIJ* on 5 mM 2-PE supplemented without Ln, with 2 µM LaCl_3_, or with 1 µM La_2_O_3_ (to provide an equimolar concentration of La^3+^) and observed no difference in growth profiles based on La^3+^ source (**Fig. S14**).

To evaluate *P. putida’s* transcriptional response to La, we cultivated wild-type *P. putida* and *P. putida* Δ*pvdL* Δ*pvdD* Δ*pvdIJ* on 2-PE without La supplementation and with 2 µM LaCl_3_ or 1 µM La_2_O_3_. Cell pellets were sampled for RNA-Seq transcriptomic analysis in late-exponential growth when 2-PE had been consumed but not fully metabolized (**Figs. 4A & S15A**). Principal component analysis indicated strong agreement between biological replicates and tight clustering between LaCl_3_ and La_2_O_3_ conditions for both strains (**Fig. 4B**). Only 91 differentially expressed genes (DEG) were observed when comparing wild-type *P. putida* on 1 µM La_2_O_3_ to 2 µM LaCl_3_ (log_2_ fold-change ≥ 1 and adjusted *p*-value ≥ 0.05, **Fig. 4C, Supplementary Excel 1**). These data support minimal transcriptional-level differentiation between La-chloride and La-oxide forms.

**Figure 4.**
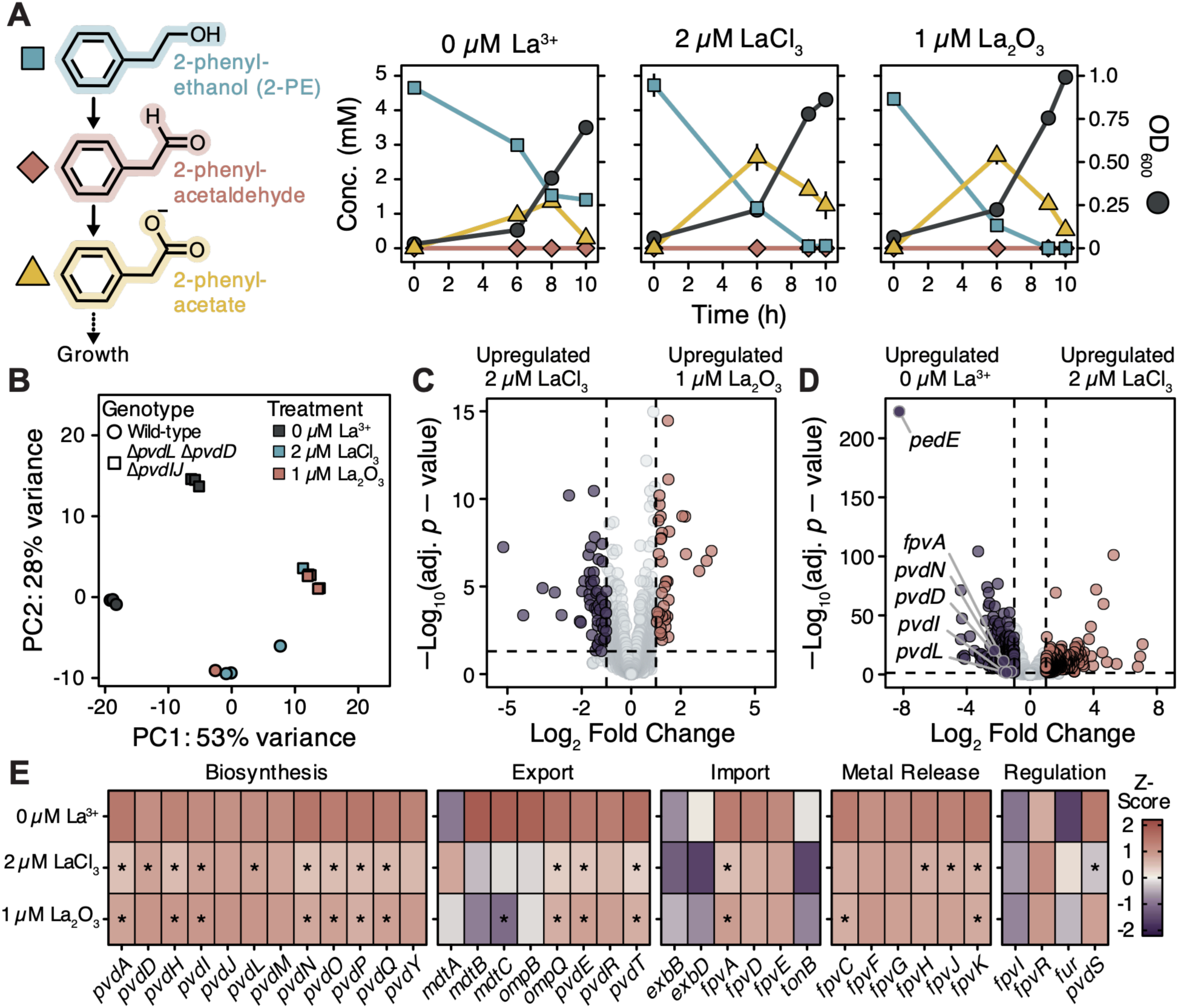
RNA-Seq transcriptomics of wild-type *P. putida* and *P. putida* a *pvdLIJD* deletion mutant on varying La supplementations. (**A**) Growth and metabolite (2-PE, 2-phenylaldehyde, and 2-phenylacetate; structures shown on left) concentrations for wild-type *P. putida* in modified MP media supplemented with 5 mM 2-PE and no La, 2 µM LaCl_3_, or 1 µM La_2_O_3_. Cultures were grown in 60 mL modified MP medium in 250 mL acid washed polycarbonate flasks at 30°C and 225 rpm. Data points represent the mean values of biological triplicates and error bars correspond to standard deviations. (**B**) PCA of rlog transformed RNA-Seq results for wild-type *P. putida* and *P. putida* Δ*pvdL* Δ*pvdD ΔpvdIJ*. Volcano plot of DEGs when comparing wild-type cultivations on (**C**) 1 µM La_2_O_3_ versus 2 µM LaCl_3_ or (**D**) 2 µM LaCl_3_ versus no La. Significance cut-offs: log_2_ fold-change ≥ 1 and Benjamini– Hochberg adjusted *p*-value ≤ 0.05, indicated by vertical and horizontal dotted black lines. (**E**) Expression (Z-score normalized) of pyoverdine-associated genes in wild-type *P. putida* cultivations. Significance is annotated by stars (*) and comparisons are between 1 µM La_2_O_3_ versus no La or 2 µM LaCl_3_ versus no La. Significance cut-offs: log_2_ fold-change ≥ 1 and Benjamini–Hochberg adjusted *p*-value ≤ 0.05.

There were 850 and 630 DEGs when comparing wild-type *P. putida* transcriptomes from 2 µM LaCl_3_ or 1 µM La_2_O_3_ to the absence of La, respectively (**Figs. 4D & S15B, Supplementary Excel 1**). In agreement with pyoverdine production conditions, the *pvd* and *fpv* gene clusters were generally down regulated in La^3+^ treated conditions compared to no La^3+^ amendment but were not differentially expressed between 1 µM La_2_O_3_ and 2 µM LaCl_3_ (adj*. p*_adj._ ≤ 0.05, |log_2_FC| ≥ 1; **Fig. 4E, Supplementary Excel 1**). Within the *pvd* and *fpv* clusters, *pvdN* and *pvdA* were the most significantly down-regulated genes in La^3+^ treated conditions compared to no La^3+^ amendment (**Supplementary Excel 1**). TonB-dependent OMRs *fpvA* and *PP_3612* exhibited the strongest responses to La addition, with both being down-regulated in 2 µM LaCl_3_ or 1µM La_2_O_3_ compared to no La conditions (**Fig. 4E, Supplementary Excel 1**). Interestingly, *PP_4222* which resides between *pvdH* and *pvdI* within the *pvd* cluster, was down-regulated upon La^3+^ addition (-2.05 and - 1.3 log_2_FC in 2 µM LaCl_3_ and 1 µM La_2_O_3_, respectively, **Supplementary Excel 1**), and encodes a homolog to the aspartyl hydroxylase SyrP protein in *Pseudomonas syringae* (*57*).

Of 38 outer membrane porins, which could mediate non-selective passage of large cations across *P. putida’s* outer membrane (*37*), 15 were found to be significantly differentially expressed in response to La supplementation (**Fig. S16, Supplementary Excel 1**). Of the significant porins, only the nonspecific porin *oprF* has been shown to allow cation diffusion across the outer membrane and was significantly down-regulated in the presence of LaCl_3_ compared to no La^3+^ (-1.26 log_2_FC, **Fig. S16, Supplementary Excel 1**) (*58*, *59*).

### La readily precipitates in LaCl_3_-supplemented media, with phenotypic changes only observed between 2-20 nM LaCl_3_

Given the similar growth and transcriptome profiles between LaCl_3_ and La_2_O_3_, we considered whether a difference in soluble La^3+^ in our media between the two supplementations was supported. As mentioned previously, Lns are known to complex with phosphates in solution forming insoluble Ln-phosphate minerals with sparing solubility products 10^-25^ to 10^-27^ (*42*, *43*). Modified M9, MP, ½ Hypho, and modified MP (developed here) contain 68 mM, 3.33 mM, 16.6 mM, and 8.5 mM phosphate respectively (*32*, *33*, *36*, *37*), all well above the typical 0.01-100 µM Ln supplementations.

To measure La precipitation in LaCl_3_-supplemented modified MP media, water, modified MP, and perturbations of modified MP medium without either phosphate or sulfate were supplemented with LaCl_3_ at pH 7 and filtered to quantify soluble La^3+^ (**Fig. 5A**). In modified MP medium, the presence of phosphate resulted in the removal of 98.4% of La^3+^. Removal of phosphate but not sulfate reduced the observed precipitation, with phosphate-free modified MP having only a 0.5% reduction (**Fig. 5A**). Acidification prior to filtration, which would re-solubilize La^3+^ from LaPO_4_, reversed these trends, indicating that phosphate in the media facilitated La precipitation as LaPO_4_.

**Figure 5.**
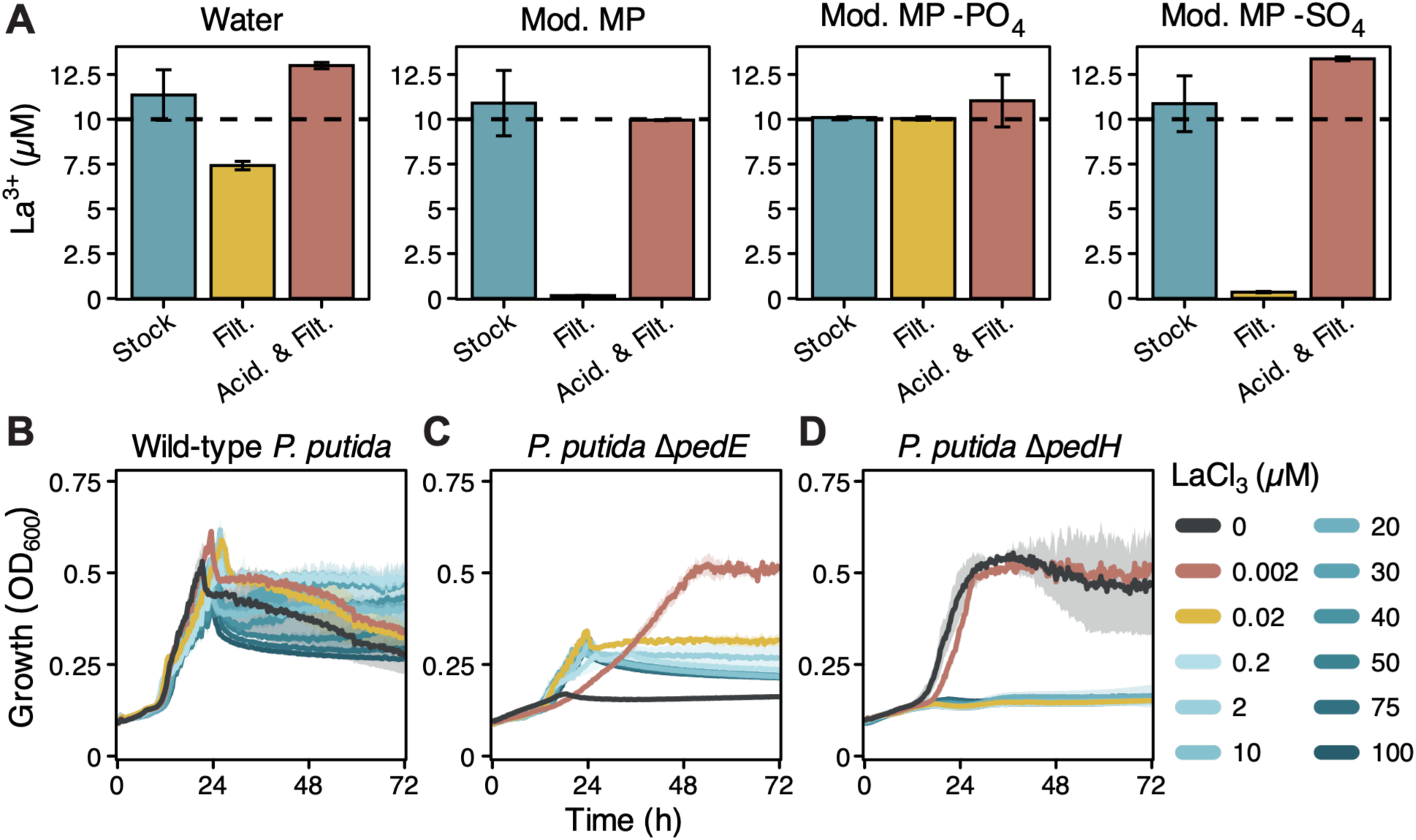
La precipitation in Modified MP medium and growth curves on 2-PE across increasing LaCl_3_ concentrations. (**A**) ICP-QQQ quantification of La in water, modified MP medium (Mod. MP), modified MP medium lacking phosphate (Mod. MP -PO_4_), and modified MP medium lacking sulfate (Mod. MP -SO_4_). Sample treatments consisted of stock solutions, 0.22 µM PES filtered solutions (Filt.), and solutions acidified to 5% final nitric acid concentration followed by 0.22 µM PES filtration (Acid. & Filt.). Bars represent the mean of values of replicate sample and error bars correspond to standard deviations. All solutions were made to a final target concentration of 10 µM La^3+^, which is indicated by the black horizontal dotted line and represents 100% recovery of La. Growth curves of (**B**) wild-type *P. putida,* (**C**) *P. putida* Δ*pedE,* and (**D**) *P. putida* Δ*pedH* in modified MP media supplemented with 0-100 μM LaCl_3_. Cultures were grown on 5 mM 2-PE in 200 µL in 100-well honeycomb plates at 30 °C. Data represent the mean values of biological triplicates and shaded regions correspond to standard deviations.

We then conducted growth assays with wild-type *P. putida* and ADH mutants with 0-100 μM LaCl_3_ supplementation (**Fig. 5; Fig. S17**) to measure phenotypic differences. PedH-dependent growth (*P. putida* Δ*pedE*) was supported at 2 nM LaCl_3_ but with markedly increased lag and max OD_600_, decreased growth rate as compared to 20 nM LaCl_3_ (**Fig. 5B**). PedE-dependent growth (*P. putida* Δ*pedH*) was permitted only at LaCl_3_ supplementations below 20 nM (**Fig. 5C**), suggesting the REE-switch is activated between 2-20 nM (pH 7, 1 µM Fe). *P. putida* Δ*pedE* growth plateaued above 200 nM, consistent with dissolved La^3+^ concentration being controlled by the K_sp_ of LaPO_4_ instead of total La^3+^ supplied. Together, this analysis suggests La-phosphates form in commonly used media formulations for Ln growth phenotyping which are bioavailable but, to some extent, insoluble.

## DISCUSSION

Based on our results, we propose a model for pyoverdine-mediated La chelation and transport across the outer membrane in *P. putida* (**Fig. 6**). In this model, the *pvdLIJD* BGC encodes an NRPS system responsible for pyoverdine and ferribactin production in the cytoplasm. While experimentally uncharacterized, these metabolites must undergo inner membrane transport and cytoplasmic pyoverdine maturation, presumably via pathways analogous to those in other pseudomonads (*60–64*), prior to extracellular export. Once exported, the metallophore chelates extracellular La, though any role in solubilization from LaPO_4_ and/or La_2_O_3_ is unknown, and the resulting complex is transported into the periplasm via FpvA-TonB-ExbBD.

**Figure 6.**
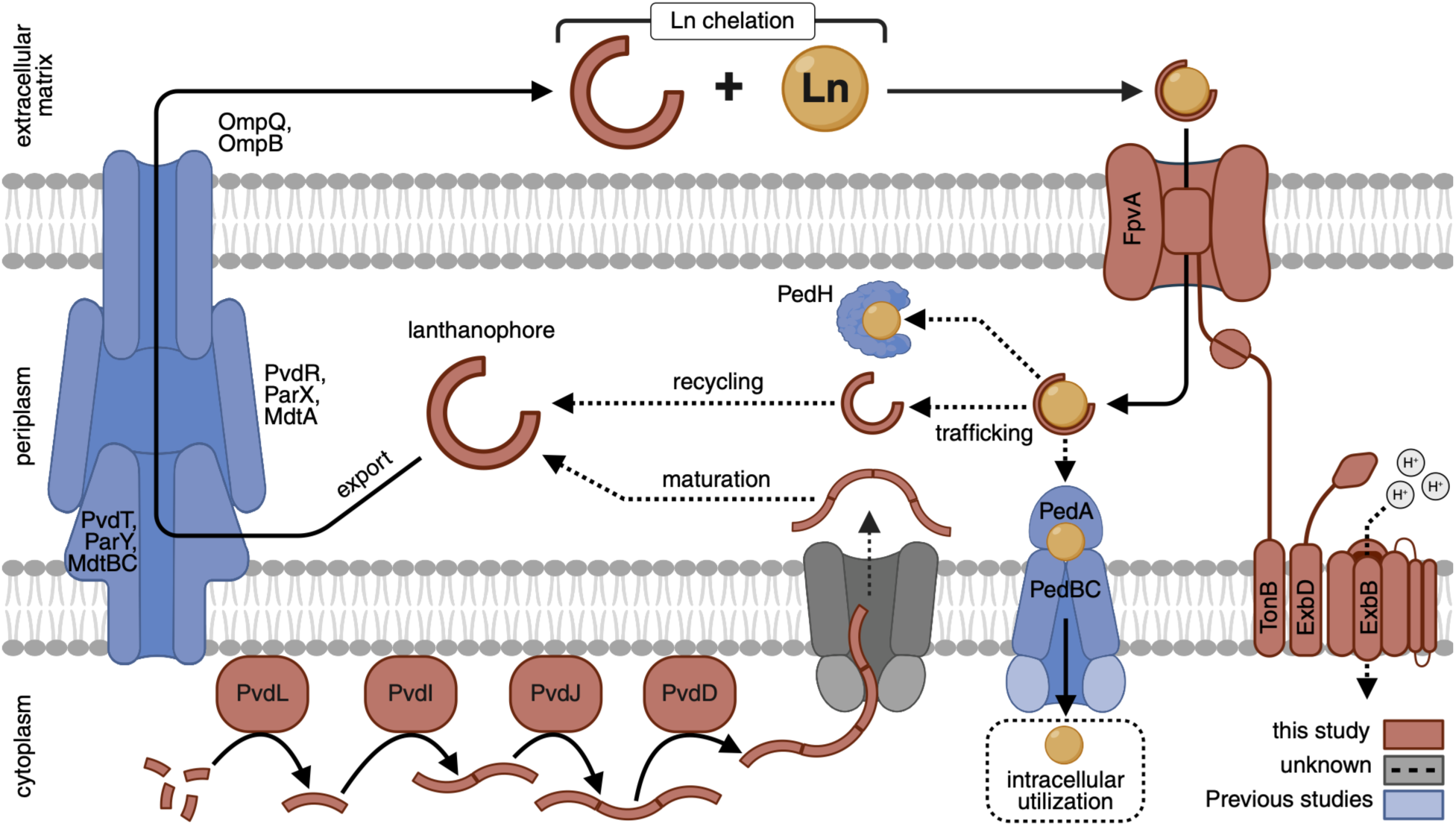
Conceptual model for pyoverdine-mediated Ln uptake in *P. putida*. Elements proposed previously are in blue (PedABC (*65*); RND export systems PvdRT-OmpQ, MdtABC-OmpB, or ParXY (*66–68*); and PedH (*69*)), and those reported here are in red (PvdLIJD, FpvA, TonB, and ExbBD). Unknown components are shown in grey and with dashed black lines. The ABC-Transporter PedA1A2BC (*PP_3338, PP_2669, PP_2849, PP_2667*) has been simplified here to PedABC for illustrative purposes.

Our results support a new functional role for bacterial pyoverdines G4R and ferribactin in Ln uptake. Pyoverdines have predominantly been studied in the context of Fe scavenging, especially within the framework of *P. aeruginosa* pathogenicity (*45*, *46*). In vitro, the hydroxamate siderophores desferrioxamine B and desferricoprogen have been shown to bind Ln^3+^ (*70*, *71*) though the physiological implications on Ln binding are unreported to our knowledge. The regulatory, biosynthetic, and chelation differences we observed between ferribactin and pyoverdine G4R are particularly intriguing. Based on homology to other pseudomonad systems, increased ferribactin abundance could be correlated with reduced acyl sidechain conversion to succinamide by the aminotransferase PvdN (*64*) and reduced chromophore maturation by the tyrosinase PvdP and/or PvdO (*60*, *61*). PP_4222 may play a similar role in metallophore maturation, as it’s homolog in *P. syringae* has been shown to convert Asp to *threo*-3-OH-Asp, a prominent residue in the pyoverdine peptide (*57*). In addition to further studies probe the biosynthetic and regulatory landscape of the maturation system, characterization of binding affinity and chelate structures with purified pyoverdine G4R and ferribactin would be highly informative in deciphering metal-binding preferences.

Worsening growth of *fpvA* and *exbBD:tonB* deletion strains with higher La supplementation may reflect increasing competition with Fe for uptake through an FpvA OMR that recognizes both metals. While the homologous FpvA-TonB-ExbBD in *P. aeruginosa* is well-characterized to facilitate Fe translocation across the outer membrane (*72*, *73*), TonB-dependent systems remain poorly characterized in *P. putida* beyond a few key studies in *P. putida* DOT-T1E implicating TonB systems in stress tolerance (*74*, *75*). Notably, however, *fpvA* was non-essential for utilization of Fe or La. These data imply redundant mechanisms for uptake of both metals, phenomena also observed for the TonB-dependent OMR LutH in an Ln-dependent *M. extorquens* strain (*76*). Prior work supports passive diffusion (*37*) or redundant and/or promiscuous OMRs (*77*, *78*), with the transcriptional profile of putative OMR *PP_3612* positioning it as a candidate. Additional outstanding questions include whether Fe- and La-chelation complexes with pyoverdine G4R and ferribactin are recognized by the same receptor, and structurally elucidation of these complexes.

The NRPS BGC was only strictly required for La-dependent growth, regardless of La form, suggesting *P. putida* employs redundant mechanisms or metallophores for Fe but not La. Metallophore redundancy is well-documented, with proposed fitness advantages including overcoming host defense systems (*79*, *80*) and “cheating” by import of externally-produced metal-chelate structures (*81*). Rhizospheric pseudomonads, known to produce an array of metallophores differing in structure and function (*77*), could similarly achieve a fitness advantage from shared metal acquisition systems, but further work is needed to understand the biogeochemical triggers for, and broader ecological relevance of, Ln scavenging, particularly in rhizospheric or phyllospheric environments.

The minimal impacts of La form (chloride versus oxide) on *P. putida’*s growth and transcriptome, compounded with poor La solubility, suggests that La^3+^ precipitates as LaPO_4_ in modified MP media. These results caution against interpreting supplemented LaCl_3_ concentrations as soluble La^3+^, where free La^3+^ would be controlled by the K_sp_ of LaPO_4_. A notable phenotypic exception was the observation that ferribactin exhibited a higher relative abundance in media with La_2_O_3_ as compared to LaCl_3_, similar to that observed for methylolanthanin (*32*). Further investigation with careful control over soluble La^3+^ (e.g., through the use of organic phosphates (*82*)) is needed to disentangle the roles of lanthanophores in solubilization versus free-ion import.

Beyond biological pathways, harnessing selective metallophores could enable critical mineral recovery or separations, including the use of engineered microbes for bioproduction or bio-inspired chemical synthesis routes. Understanding lanthanophore biosynthesis and regulatory networks could enable overproduction in engineered *P. putida,* with potential applications in bioleaching and bioremediation (*28*, *83–88*). Similarly, deciphering trafficking pathways could uncover additional small molecule chelators or Ln-binding proteins, with demonstrated applications in Ln separations (*26*, *89*, *90*).

## MATERIALS AND METHODS

### Culture conditions

Wild-type *P. putida* (ATCC 47054) was used for all experiments, or further strain construction, and overnight cultures were revived from 20% (v/v) glycerol stocks in 20 mL of Miller’s LB (Sigma #L3522) in 125 mL baffled flasks at 30 °C and 225 rpm. For preliminary growth experiments, *P. putida* was cultivated in either modified M9 with trace elements solution or MP medium with C7 metals supplementation as described previously (*33*). After initial media refinement, modified MP medium was used for all subsequent experiments and contained 8.5 mM phosphate buffer (pH 7, Na_2_HPO_4_ and NaH_2_PO_4_), 8 mM (NH₄)₂SO₄, 0.5 mM MgCl₂, 100 µM CaCl₂, 45 µM Na_3_-Citrate, 1 µM FeSO₄×7H₂O. Specifics regarding media and metal solutions may be found in **Tables S2 and S3**. 5 mM 2-phenylethanol (Sigma #77861) was used as the sole carbon source in all experiments. Cultures were amened with LaCl_3_×7H₂O (Sigma #262072) or La_2_O_3_ (Sigma #289205) as indicated. Individual stock solutions for all reagents were dissolved in MilliQ water and 50 mL polypropylene tubes, then 0.22 µM sterile filtered using the Steriflip^®^ filtration system (Millipore, USA). La_2_O_3_ was made with sterile MilliQ water and sterilized by pasteurization at 60 °C for 1 h. Solutions were made without metal or glass equipment to avoid trace contaminations of metals. Stock solutions were checked via ICP-MS to ensure no Ln contamination and La stock concentrations were similarly validated. All components were sterilely combined and aliquoted prior to inoculation.

### Strain construction

Genetic deletions were performed using homologous recombination and the antibiotic/*sacB* method for gene knockouts, as described previously (*91*). In brief, plasmids were electroporated in *P. putida* strains, two rounds of selection were conducted on LB + 50 μg/mL kanamycin (kan50) agar plates, followed by two rounds of counter selection on YT + 25% sucrose plates, then isolate colonies were patched onto both LB and LB + kan50 agar plates, and colony PCR was conducted on LB patch colonies to verify gene knockouts (**Tables S6-8**). Antibiotic and sucrose selection places were amended with 20 µM FeSO₄×7H₂O and 20 µM ferric ammonium citrate for Δ*fpvA* and Δ*exbBD:tonB* knockouts to supplement decreased Fe uptake ability. For colony PCR, colonies directly from LB patch plates were picked with a sterile pipette tip into 20 µL of PCR reaction mixture. Reaction mixtures consisted of 1× MyTaq HS Red master mix (Bioline, USA), 0.5 µM forward primer, 0.5 µM reverse primer, 7 µL PCR grade water, 1 µL DMSO (5% final concentration). PCRs were conducted with an initial boil at 95 °C for 5 min followed by 30 cycles of 95 °C for 30 s, 68 °C for 30 s, and 72 °C for a variable time to amplify desired amplicons assuming 30 s/kb, and lastly a final extension at 72 °C for 2 min and 4 °C hold. Confirmed knockouts were incubated in LB overnight and cryostocked at 20% glycerol and -80 °C until further experimentation. Plasmids were ordered from Twist Bioscience (San Francisco, USA) with 1 kilobase up and downstream homology regions in the pK18sB vector backbone containing *sacB* and *kanR* (**Table S8**). Sequenced genomes for *P. putida* Δ*pedE* and *P. putida* Δ*pvdL* Δ*pvdD* Δ*pvdIJ* mutants can be found at the National Center for Biotechnology Information (NCBI) Sequence Read Archive (SRA) under Bioproject PRJNA1512547.

### Growth experiments

Growth phenotyping was conducted in biological triplicate in either 100-well honeycomb plates in a BioscreenC™ (Growth Curves USA, USA) or baffled shake flasks. Sterile controls consisting of prepared media inoculated with wash solution were always conducted in parallel. BioscreenC™ experiments were conducted with 200 µL modified M9, MP, or modified MP medium at 30 °C with maximum continuous shaking and OD_600_ measurements occurring every 15 min. Cells from overnight cultures were prepared for inoculations by centrifugation at 4,500×*g* for 7 min followed by removal of supernatant. Next, cells underwent two rounds of resuspension in 50% of the initial volume with wash solution, centrifugation at 4,500×*g* for 7 min, and removal of supernatant. Cells were then resuspended with wash solution to 20% of the initial volume. OD_600_ was then quantified, and cells were diluted to a working OD_600_ of 5. All experiments were inoculated to an initial OD_600_ of 0.05. During testing, we discovered that cell wash solution for inoculation impacts growth viability (**Table S9**, **Fig. S18A**), and it was determined that including 30 mM PIPES in the cell wash solution was necessary for growth (**Fig. S18B**).

Flask experiments were conducted in 125 mL and 250 mL baffled polycarbonate flasks (Nest Scientific, USA). Flasks were acid washed with 4 N HCl to remove any residual metals and autoclaved prior to use. Incubations were conducted at 30 °C and 225 rpm. OD_600_ measurements were conducted via spectrophotometer measurements. Samples were taken for transcriptomic, 2-phenylethanol, and metabolomics analysis as indicated (described below). Lag phase max growth rate calculations were performed using Welly and the tangent intercept method (*92*). Growth curves were constructed in R using the ggplot2 plotting package (*93*, *94*).

Cultures for RNA sequencing, 2-phenylethanol quantification, and 2-PE analysis were conducted in 60 mL modified MP medium supplemented with either 0 µM La, 2 µM LaCl_3_, or 1 µM La_2_O_3_ in 250 mL flasks. For metabolite quantization, culture supernatant was centrifuged at 15,000×*g* for 2 min at 4 °C, 0.22 µM PES filtered, 1:2 diluted with MilliQ water in amber HPLC vials, capped and stored at -20 °C until instrument analysis.

For pyoverdine analysis, 47 mL of late-exponential phase cultures were transferred to 50 mL polypropylene tubes and centrifuged at 4,500×*g* for 10 min at 4 °C. 45 mL of supernatant was transferred to a fresh polypropylene tube, flash frozen in liquid nitrogen, and stored at -80 °C until metabolite extraction and purification.

### Transcriptomic analysis

Cultures were grown to late-exponential phase, and 1.5 mL culture was collected and centrifuged at 15,000×*g* and 4 °C for 3 min to harvest cells. Next, supernatant was removed and pellets were flash frozen in liquid nitrogen then stored at -80 °C until shipment. Cell pellets were shipped on dry ice to Genewiz for sequencings (Plainfield, USA). Genewiz performed their standard RNA-Seq pipeline, including RNA extraction, rRNA depletion, library preparation, and Illumina® NovaSeq™ 2×150 bp sequencing with a Q-score cut-off of >30. Raw FASTQ files were processed in Kbase (*95*), where read quality was assessed using FastQC version 0.12.1 (*96*), trimmed using Trimmomatic version 0.39 (*97*), aligned to the *P. putida* KT2440 reference genome using Bowtie2 version 2.3.2 (*98*), and assembled with StringTie version 2.1.5 (*99*). Transcript counts were imported into R where differential gene expression analysis was conducted using DESeq2 version 1.20.0 (*100*). Heatmaps depicting transcript abundance data were first processes using dds() then normalized using rlog(), both functions in DESeq2, Z-scores were calculated by gene. Raw sequencing reads for RNA-Seq analysis may be found at the NCBI SRA under BioProject PRJNA1512547.

### Quantification of lanthanides

Ln series elements, omitting Promethium (Pm), were quantified using an Agilent 8900 triple-quadrupole inductively coupled plasma mass spectrometer (ICP-QQQ) equipped with an Ultra High Matrix Introduction (UHMI) system, a MicroMist nebulizer, and nickel-plated interface cones to provide enhanced matrix tolerance, robustness, and corrosion resistance. The instrument was operated in helium (He) mode using HMI-8 plasma mode with preconfigured parameters (X-Lens optics, 7 replicates with 100 sweeps per replicate, 70 s sample uptake time, 50 s stabilization time, and a nebulizer pump speed of 0.2 rps), and polyatomic interferences were removed through kinetic energy discrimination (KED) within the collision/reaction cell (CRC) and a gas flow of 5.5 mL/min. Samples were diluted 2-5X as needed, and all samples and calibration standards were acidified to 3.5% HNO_3_ using parts-per-trillion (ppt) grade acid (GFS chemicals Cat.# 72030) to ensure low background levels and were introduced into the system using a SPS 4 autosampler.

Quality control (QC) checks, including blanks and calibration-verification standards (CVS), were run at least every 20 samples. The intermediate calibration standard used to prepare the calibration curve was prepared by mixing 1.0 mL of Rare-Earth Element ICP mix (Sigma-Aldrich Cat. ##67349-100 mL), 0.5 mL of transition elements ICP standard (100 ppm; Inorganic Ventures Cat.# CCS-6), and 5.0 mL of the ICP aluminum standard (1000 µg/mL; InorganicVentures Cat.# AAAL1 with 43.5 mL of 3.5% nitric acid. The resulting intermediate standard contained all target elements at a concentration of 1000 ng/mL, and was used to generate a 13-point curve (0 to 250.000 ppb) using a blank-offset origin with no weighting. Correlation coefficients for all analytes was ≥ 0.995. Blanks were required to fall below the lower quantitation limit (LQL), and CVS recoveries were accepted if results were within 85–115% of the expected concentration. Additionally, for results to be acceptable they must maintain an acceptable RSD of counts per second between replicates, in this case a threshold of ±10%, and demonstrated stable internal standard behavior.

Internal standards were used to correct for plasma-matrix interactions, signal suppression, and instrument drift. Four AccuStandard stock solutions were used as internal standards: indium (ICP-MS-25N-0.01X-1), rhodium (ICP-MS-46H-0.01X-1), bismuth (ICP-MS-06N-0.01X-1), and germanium (ICP-21W-1). To prepare the 500 ng/mL intermediate internal standard, 1.25 mL each of the indium, rhodium, and bismuth standards and 0.125 mL of the germanium standard were combined in a 250 mL poly-bottle and brought to a final volume of 250 mL with 3.5% nitric acid. Internal standard recoveries were required to remain within 50–150%.

### Pyoverdine/ferribactin identification and molecular networking

Solid phase extraction was used to obtain metabolites from 22.5 mL culture supernatant using Chromabond HLB (Macherey-Nagel) columns equilibrated with 3 column volumes of methanol (LCMS grade, Fisher Chemical) followed by 3 column volumes of water (LCMS grade, Fisher Chemical).

Methanol (100%) was used for elution in 1.5 mL, which was then dried overnight using a centrivap (Eppendorf). Samples were resuspended in 50% methanol/water to reach 5 mg/mL, or 100 µg/mL for untargeted analysis.

Reverse-phase chromatography was run on a Vanquish UHPLC system coupled to a Q-Exactive HF quadrupole orbitrap (Thermo Scientific). A 1.7 µm C18 EVO column was used, with dimensions 150 x 2.1 mm (Phenomenex, #00F-4726-AN), at 0.5 mL/min flow rate and 25 °C column temperature. The mobile phase was water with 0.1% formic acid (A) and acetonitrile with 0.1% formic acid (B), and 5 μL of sample was injected and eluted using the following gradient: 0-9 min, 2-50% B; 9-10 min, 50-99% B; followed by a 3 min washout phase at 99% B and a 3 min re-equilibration phase at 2% B.

Data dependent (DDA) of MS/MS was performed in positive mode with technical triplicates for each biological triplicate. The MS scan range was set to 150– 1500 m/z with a resolution of m/z 200 of 120,000 with one microscan. The maximum ion injection time was set to 100 ms with an automated gain control (AGC) target of 1.0E6. Up to five MS/MS spectra per MS1 survey scan were recorded in DDA mode with resolution m/z 200 of 15,000 with one micro-scan. The maximum ion injection time for MS/MS scans was set to 200 ms with an AGC target of 5E5 ions. Normalized collision energy was set to a stepwise increase from 25 to 35 to 45%. MS/MS scans were triggered at the apex of chromatographic peaks within 2–15 s from their first occurrence. Dynamic precursor exclusion was set to 5 s. Quality control (QC) mixes, method blanks, and instrument blanks were analyzed alongside the samples to ensure validity of analysis.

The raw organic MS files were centroided to .mzML using MSConvert (*101*) and untargeted analysis was performed using mzmine 4.9.14 (*102*), GNPS2 (*103*) and the associated FBMN-STATS GUI (*104*), and R version 4.4.2 (*93*). Converted .mzML files were subjected to feature detection in mzmine using the parameters specified in the batch file (see *Data Availability*), including: MS1 mass detection of 5E3, MS2 mass detection noise factor 2.5, chromatogram building between retention time 0.25 to 10 min., join aligner m/z tolerance sample to sample 5 ppm, and retention time tolerance 0.20 min. The feature quantitation table (.csv), MS2 spectra (.mgf), and sample metadata (.csv) were uploaded to the Global Natural Product Social 2 (GNPS2) molecular networking platform for feature based molecular networking. After feature finding, media blanks were subtracted in the Hitchhiker’s Guide with a 0.3 cutoff and samples were normalized to TIC. Imputation was performed, although non-imputed data were used for downstream analyses involving presence/absence metrics and to quantify metabolite presence across sample groups. GNPS2 was used for molecular networking and MassQL queries were performed using the tool present in the GNPS2 network browser (see *Data Availability*).

### Metal-binding via metal-infusion mass spectrometry

For metal infusions, a 1.7 µm C18 EVO column (50 x 1.0 mm; Phenomenex, #00B-4726-A0), 0.15 mL/min flow rate, and 25 °C column temperature with post-column metal infusion loop was used, as described previously (*54*). Briefly, flow from the column was combined with inflow from the quaternary pump for pH adjustment (0.15 mL/min of 10 mM ammonium acetate) using a T-connector such that the outflow was perpendicular to both inflows. Metal was infused using a syringe pump (Chemyx F100T2) and Hamilton syringe (Hamilton 81265) flowing at 5 µL/min, with influent metal solution meeting the pH-adjusted column solution perpendicularly. The Hamilton syringe was equipped with a PEEK needle (Hamilton 8650-01) which was equipped with a finger-tight fitting for a direct connection to the sample loop. LaCl_3_ was infused at 3 mM in pure LCMS water (Fisher Chemical), while La/Fe (Fisher Chemical) combination infusions were performed with 1 mM of each metal in water. Metal solutions were prepared fresh on the day of each infusion experiment. For metal infusions, a 1 mg/mL solution of each supernatant extract was used, and 5 μL of sample was injected and eluted using the following gradient elution: 0-8 min, 2-50% B, 8-10 min, 50-99% B followed by a 2 min washout phase at 99% B and a 4 min re-equilibration phase at 2% B.

The MS scan range was 250-1500 m/z to avoid the infused LaCl_3_ mass of 243.8129, and MS analysis was performed in positive mode with heated electrospray ionization (HESI) parameters were set to 35 L/min sheath gas flow, 10 L/min auxiliary gas flow. The spray voltage was set to 3.5 kV and the inlet capillary to 250 °C and 55 V S-lens level was applied. A resolution of m/z 200 of 45,000 with one microscan was set. The maximum ion injection time was set to 100 ms with an automated gain control (AGC) target of 1.0E6. DDA of MS2 spectra was performed with the parameters listed above but MS2 information was not used in analysis of metal binding. Raw files and a chart of peak quantifications can be found on MASSIVE (see *Data Availability*). For known metal binding quantification, Qualbrowser (Xcalibur version 4.6.67.17) was used to identify base peaks and acquire peak areas to assess metal binding. Peaks were quantified using automatic peak detection, with manual peak annotation if automatic peak detection failed to capture the full peak. Peak detection was performed with a baseline window of 40, area noise factor of 5, and peak noise factor of 10.

### Quantification of phenylacetaldehyde, phenylacetic acid, and 2-phenylethanol

Phenylacetaldehyde, phenylacetic acid, and 2-phenylethanol were quantified by reverse-phase ultra-high-performance liquid chromatography on an Agilent 1290 Infinity II system (Agilent Technologies) equipped with a diode array detector (DAD). Separation of the analytes was achieved with an Atlantis Premier BEH C18 AX column (1.7 µm, 2.1 × 100 mm; Waters) held at 40.0 °C. The following gradient was employed: 0.2% formic acid in water (A) and acetonitrile (B) using a flowrate of 0.600 mL/min, increasing from 10% to 99% B over 4.00 min before returning to initial conditions for re-equilibration with a total run time of 5.50 min. Samples and standards were injected at a volume of 2.00 µL. Analytes were detected and quantified by DAD at a wavelength of 265 nm. External calibration curves for each analyte were produced at a range of 5–1000 ppm using a linear regression fit, with R^2^ ≥ 0.995 for all three compounds.

## Acknowledgements

This work was authored in part by the National Laboratory of the Rockies for the U.S. Department of Energy (DOE), operated under Contract No. DE-AC36-08GO28308. K.R., A.B., K.R., and M.I. were supported in-part by the U.S. DOE, Office of Science, Biological and Environmental Research program.

J.M. and A.T.A. were supported by the National Science Foundation under Award Number CHE-2441115.

K.R. and A.Z.W. were additionally supported by the Laboratory Directed Research and Development (LDRD) Program at the National Laboratory of the Rockies. The views expressed in the article do not necessarily represent the views of the DOE or the U.S. Government. The U.S. Government retains a nonexclusive, paid-up, irrevocable, worldwide license to publish or reproduce the published form of this work, or allow others to do so, for U.S. Government purposes.

## Author Contributions

K.R., A.T.A., and A.Z.W. conceptualized the work. K.R. constructed stains, performed bacterial growth experiments, and conducted RNA-sequencing. J.M. and A.T.A. performed pyoverdine metabolomics and metal-infusion MS experiments. A.B., K.R., and M.I. performed metal and metabolite quantitation. K.R., J.M., A.T.A., and A.Z.W wrote the manuscript with contribution from all authors. A.T.A. and A.Z.W. provided supervision and funding acquisition.

## Data Availability

All numerical data presented in the main text is provided as in Supplementary Excel 1. Unprocessed fastq files from RNA-sequencing and sequenced whole-genomes for selected strains were deposited in the NCBI SRA under BioProject ID RJNA1512547. The feature based molecular network can be found at: https://gnps2.org/status?task=97333032e0e9455ebbdc35388b131224. MS data files, including mzmine batch files containing parameters, were uploaded to MASSIVE under the IDs MSV000102931 and MSV000102969.

## References

1. R. R. Crichton, “Chapter 1: An overview of the role of metals in biology” in Practical Approaches to Biological Inorganic Chemistry (Second *Edition)*, R. R. Crichton, R. O. Louro, Eds. (Elsevier, 2020), pp. 1–16.

2. C. Andreini, I. Bertini, G. Cavallaro, G. L. Holliday, J. M. Thornton, Metal ions in biological catalysis: from enzyme databases to general principles. J Biol Inorg Chem (2008).

3. M. Kobayashi, S. Shimizu, Cobalt proteins. Eur. J. Biochem. 261, 1–9 (1999).

4. R. Hansch, R. R. Mendel, Physiological functions of mineral micronutrients (Cu, Zn, Mn, Fe, Ni, Mo, B, Cl). Curr. Opin. Plant Biol. 12, 259–266 (2009).

5. E. I. Solomon, U. M. Sundaram, T. E. Machonkin, Multicopper Oxidases and Oxygenases. Chem. Rev. 96, 2563–2600 (1996).

6. T. Dudev, C. Lim, Competition among Metal Ions for Protein Binding Sites: Determinants of Metal Ion Selectivity in Proteins. Chem. Rev. 114, 538–556 (2014).

7. H. Irving, R. J. P. Williams, Order of Stability of Metal Complexes. Nature 162, 746–747 (1948).

8. J. D. Helmann, Metals in Motion: Understanding Labile Metal Pools in Bacteria. Biochemistry 64, 329–345 (2025).

9. S. Gama, R. Hermenau, M. Frontauria, D. Milea, S. Sammartano, C. Hertweck, W. Plass, Iron Coordination Properties of Gramibactin as Model for the New Class of Diazeniumdiolate Based Siderophores. Chem. – Eur. J. 27, 2724–2733 (2021).

10. S. M. Kraemer, O. W. Duckworth, J. M. Harrington, W. D. C. Schenkeveld, Metallophores and Trace Metal Biogeochemistry. Aquat. Geochem. 21, 159–195 (2015).

11. J. Kramer, Ö. Özkaya, R. Kümmerli, Bacterial siderophores in community and host interactions. Nat. Rev. Microbiol. 18, 152–163 (2020).

12. R. C. Hider, X. Kong, Chemistry and biology of siderophores. Nat. Prod. Rep. 27, 637 (2010).

13. M. Knapp, L.-A. Giddings, Bacterial Siderophore Production in Metal-Rich Environments: Underexplored Sources of Siderophores and Insights into Bioremediation. J. Nat. Prod. 89, 846–863 (2026).

14. R. Soares, I. B. Trindade, R. O. Louro, Iron binding before iron limitation: siderophore synthesis arose well before iron became a limiting element. ISME Commun. 6, ycag025 (2026).

15. J. H. Crosa, C. T. Walsh, Genetics and Assembly Line Enzymology of Siderophore Biosynthesis in Bacteria. Microbiol. Mol. Biol. Rev. 66, 223–249 (2002).

16. A. Silale, B. Van Den Berg, TonB-Dependent Transport Across the Bacterial Outer Membrane. Annu. Rev. Microbiol. 77, 67–88 (2023).

17. N. Noinaj, M. Guillier, T. J. Barnard, S. K. Buchanan, TonB-Dependent Transporters: Regulation, Structure, and Function. Annu. Rev. Microbiol. 64, 43–60 (2010).

18. G. Ganne, K. Brillet, B. Basta, B. Roche, F. Hoegy, V. Gasser, I. J. Schalk, Iron Release from the Siderophore Pyoverdine in *Pseudomonas aeruginosa* Involves Three New Actors: FpvC, FpvG, and FpvH. ACS Chem. Biol. 12, 1056–1065 (2017).

19. I. J. Schalk, L. Guillon, Fate of ferrisiderophores after import across bacterial outer membranes: different iron release strategies are observed in the cytoplasm or periplasm depending on the siderophore pathways. Amino Acids 44, 1267–1277 (2013).

20. P. Delepelaire, Bacterial ABC transporters of iron containing compounds. Res. Microbiol. 170, 345–357 (2019).

21. M. Aide, “Lanthanide Soil Chemistry and Its Importance in Understanding Soil Pathways: Mobility, Plant Uptake, and Soil Health” in Lanthanides (IntechOpen, 2018; https://www.intechopen.com/chapters/62330).

22. Y. Hibi, K. Asai, H. Arafuka, M. Hamajima, T. Iwama, K. Kawai, Molecular structure of La3+-induced methanol dehydrogenase-like protein in *Methylobacterium radiotolerans*. J. Biosci. Bioeng. 111, 547–549 (2011).

23. J. A. Cotruvo, Jr., E. R. Featherston, J. A. Mattocks, J. V. Ho, T. N. Laremore, Lanmodulin: A Highly Selective Lanthanide-Binding Protein from a Lanthanide-Utilizing Bacterium. J. Am. Chem. Soc. 140, 15056–15061 (2018).

24. W. B. Larrinaga, J. J. Jung, C.-Y. Lin, A. K. Boal, J. A. Cotruvo, Modulating metal-centered dimerization of a lanthanide chaperone protein for separation of light lanthanides. Proc. Natl. Acad. Sci. 121, e2410926121 (2024).

25. J. L. Hemmann, P. Keller, L. Hemmerle, T. Vonderach, A. M. Ochsner, M. Bortfeld-Miller, D. Günther, J. A. Vorholt, Lanpepsy is a novel lanthanide-binding protein involved in the lanthanide response of the obligate methylotroph *Methylobacillus flagellatus*. J. Biol. Chem. 299, 102940 (2023).

26. Z. Dong, J. A. Mattocks, G. J.-P. Deblonde, D. Hu, Y. Jiao, J. A. Jr. Cotruvo, D. M. Park, Bridging Hydrometallurgy and Biochemistry: A Protein-Based Process for Recovery and Separation of Rare Earth Elements. ACS Cent. Sci. 7, 1798–1808 (2021).

27. Z. Dong, J. A. Mattocks, J. A. Seidel, J. A. Cotruvo, D. M. Park, Protein-based approach for high-purity Sc, Y, and grouped lanthanide separation. Sep. Purif. Technol. 333, 125919 (2024).

28. H. S. Zurier, S. Banta, D. M. Park, D. W. Reed, A. Z. Werner, Biotechnological solutions for critical mineral recovery from unconventional feedstocks. Curr. Opin. Biotechnol. 95, 103336 (2025).

29. J. A. Cotruvo, The Chemistry of Lanthanides in Biology: Recent Discoveries, Emerging Principles, and Technological Applications. ACS Cent. Sci. 5, 1496–1506 (2019).

30. E. R. Featherston, J. A. Cotruvo, The biochemistry of lanthanide acquisition, trafficking, and utilization. Biochim. Biophys. Acta BBA - Mol. Cell Res. 1868, 118864 (2021).

31. P. O. Juma, Y. Fujitani, O. Alessa, T. Oyama, H. Yurimoto, Y. Sakai, A. Tani, Siderophore for Lanthanide and Iron Uptake for Methylotrophy and Plant Growth Promotion in Methylobacterium aquaticum Strain 22A. Front. Microbiol. 13, 921635 (2022).

32. A. M. Zytnick, S. M. Gutenthaler-Tietze, A. T. Aron, Z. L. Reitz, M. T. Phi, N. M. Good, D. Petras, L. J. Daumann, N. C. Martinez-Gomez, Identification and characterization of a small-molecule metallophore involved in lanthanide metabolism. Proc. Natl. Acad. Sci. 121, e2322096121 (2024).

33. M. Wehrmann, P. Billard, A. Martin-Meriadec, A. Zegeye, J. Klebensberger, Functional Role of Lanthanides in Enzymatic Activity and Transcriptional Regulation of Pyrroloquinoline QuinoneDependent Alcohol Dehydrogenases in Pseudomonas putida KT2440. mBio 8, e00570–17 (2017).

34. M. G. Thompson, M. R. Incha, A. N. Pearson, M. Schmidt, W. A. Sharpless, C. B. Eiben, P. Cruz-Morales, J. M. Blake-Hedges, Y. Liu, C. A. Adams, R. W. Haushalter, R. N. Krishna, P. Lichtner, L. M. Blank, A. Mukhopadhyay, A. M. Deutschbauer, P. M. Shih, J. D. Keasling, Fatty Acid and Alcohol Metabolism in *Pseudomonas putida*: Functional Analysis Using Random Barcode Transposon Sequencing. Appl. Environ. Microbiol. 86, e01665–20 (2020).

35. B. Mückschel, O. Simon, J. Klebensberger, N. Graf, B. Rosche, J. Altenbuchner, J. Pfannstiel, A. Huber, B. Hauer, Ethylene glycol metabolism by *Pseudomonas putida*. Appl. Environ. Microbiol. 78, 8531–8539 (2012).

36. M. Wehrmann, C. Berthelot, P. Billard, J. Klebensberger, The PedS2/PedR2 Two-Component System Is Crucial for the Rare Earth Element Switch in Pseudomonas putida KT2440. mSphere 3, e00376–18 (2018).

37. M. Wehrmann, C. Berthelot, P. Billard, J. Klebensberger, Rare Earth Element (REE)-Dependent Growth of Pseudomonas putida KT2440 Relies on the ABC-Transporter PedA1A2BC and Is Influenced by Iron Availability. Front. Microbiol. 10 (2019).

38. V. de Lorenzo, D. Pérez-Pantoja, P. I. Nikel, *Pseudomonas putida* KT2440: the long journey of a soil-dweller to become a synthetic biology chassis. J. Bacteriol. 206, e0013624 (2024).

39. P. I. Nikel, V. de Lorenzo, *Pseudomonas putida* as a functional *chassis* for industrial biocatalysis: From native biochemistry to *trans*-metabolism. Metab. Eng. 50, 142–155 (2018).

40. R. R. Klauer, D. A. Hansen, D. Wu, L. M. O. Monteiro, K. V. Solomon, M. A. Blenner, Biological Upcycling of Plastics Waste. Annu. Rev. Chem. Biomol. Eng. 15, 315–342 (2024).

41. H.-W. Zhu, C. Wang, H.-Y. Jia, Z.-H. Liu, B.-Z. Li, Engineered *Pseudomonas putida* : a versatile chassis for lignin valorization. Green Chem. 27, 10316–10345 (2025).

42. F. H. Firsching, S. N. Brune, Solubility products of the trivalent rare-earth phosphates. J. Chem. Eng. Data 36, 93–95 (1991).

43. R. Kijkowska, R. Z. LeGeros, Preparation and Properties of Lanthanide Phosphates. Key Eng. Mater. 284–286, 79–82 (2005).

44. K. Blin, S. Shaw, H. E. Augustijn, Z. L. Reitz, F. Biermann, M. Alanjary, A. Fetter, B. R. Terlouw, W. W. Metcalf, E. J. N. Helfrich, G. P. van Wezel, M. H. Medema, T. Weber, antiSMASH 7.0: new and improved predictions for detection, regulation, chemical structures and visualisation. Nucleic Acids Res. 51, W46–W50 (2023).

45. I. J. Schalk, L. Guillon, Pyoverdine biosynthesis and secretion in Pseudomonas aeruginosa: implications for metal homeostasis. Environ. Microbiol. 15, 1661–1673 (2013).

46. M. T. Ringel, T. Brüser, The biosynthesis of pyoverdines. Microb. Cell 5, 424–437 (2018).

47. A. L. M. Salah El Din, P. Kyslík, D. Stephan, M. A. Abdallah, Bacterial iron transport: Structure elucidation by FAB-MS and by 2D NMR (1H, 13C, 15N) of pyoverdin G4R, a peptidic siderophore produced by a nitrogen-fixing strain of Pseudomonas putida. Tetrahedron 53, 12539–12552 (1997).

48. M. Baune, Y. Qi, D. A. Volmer, H. Hayen, Structural characterization of pyoverdines produced by Pseudomonas putida KT2440 and Pseudomonas taiwanensis VLB120. BioMetals 30, 589–597 (2017).

49. H. Wei, L. Aristilde, Structural characterization of multiple pyoverdines secreted by two Pseudomonas strains using liquid chromatography-high resolution tandem mass spectrometry with varying dissociation energies. Anal. Bioanal. Chem. 407, 4629–4638 (2015).

50. K. Rehm, V. Vollenweider, R. Kümmerli, L. Bigler, Rapid identification of pyoverdines of fluorescent Pseudomonas spp. by UHPLC-IM-MS. BioMetals 36, 19–34 (2022).

51. Y. S. Cody, D. C. Gross, Characterization of Pyoverdin_pss_ , the Fluorescent Siderophore Produced by *Pseudomonas syringae* pv. *syringae*. Appl. Environ. Microbiol. 53, 928–934 (1987).

52. T. Damiani, A. K. Jarmusch, A. T. Aron, D. Petras, V. V. Phelan, H. N. Zhao, W. Bittremieux, D. D. Acharya, M. M. A. Ahmed, A. Bauermeister, M. J. Bertin, P. D. Boudreau, R. M. Borges, B. P. Bowen, C. J. Brown, F. O. Chagas, K. D. Clevenger, M. S. P. Correia, W. J. Crandall, M. Crüsemann, E. Fahy, O. Fiehn, N. Garg, W. H. Gerwick, J. R. Gilbert, D. Globisch, P. W. P. Gomes, S. Heuckeroth, C. A. James, S. A. Jarmusch, S. A. Kakhkhorov, K. B. Kang, N. Kessler, R. D. Kersten, H. Kim, R. D. Kirk, O. Kohlbacher, E. E. Kontou, K. Liu, I. Lizama-Chamu, G. T. Luu, T. Luzzatto Knaan, H. Mannochio-Russo, M. T. Marty, Y. Matsuzawa, A. C. McAvoy, L.-I. McCall, O. G. Mohamed, O. Nahor, H. Neuweger, T. H. J. Niedermeyer, K. Nishida, T. R. Northen, K. E. Overdahl, J. Rainer, R. Reher, E. Rodriguez, T. T. Sachsenberg, L. M. Sanchez, R. Schmid, C. Stevens, S. Subramaniam, Z. Tian, A. Tripathi, H. Tsugawa, J. J. J. van der Hooft, A. Vicini, A. Walter, T. Weber, Q. Xiong, T. Xu, T. Pluskal, P. C. Dorrestein, M. Wang, A universal language for finding mass spectrometry data patterns. Nat. Methods 22, 1247–1254 (2025).

53. K. Rehm, V. Vollenweider, R. Kümmerli, L. Bigler, A comprehensive method to elucidate pyoverdines produced by fluorescent Pseudomonas spp. by UHPLC-HR-MS/MS. Anal. Bioanal. Chem. 414, 2671–2685 (2022).

54. A. T. Aron, D. Petras, R. Schmid, J. M. Gauglitz, I. Büttel, L. Antelo, H. Zhi, S.-P. Nuccio, C. C. Saak, K. P. Malarney, E. Thines, R. J. Dutton, L. I. Aluwihare, M. Raffatellu, P. C. Dorrestein, Native Mass Spectrometry based Metabolomics Identifies Metal-binding Compounds. Nat. Chem. 14, 100–109 (2022).

55. J. M. Rand, T. Pisithkul, R. L. Clark, J. M. Thiede, C. R. Mehrer, D. E. Agnew, C. E. Campbell, A. L. Markley, M. N. Price, J. Ray, K. M. Wetmore, Y. Suh, A. P. Arkin, A. M. Deutschbauer, D. Amador-Noguez, B. F. Pfleger, A metabolic pathway for catabolizing levulinic acid in bacteria. Nat. Microbiol. 2, 1624–1634 (2017).

56. D. Cobessi, H. Celia, N. Folschweiller, I. J. Schalk, M. A. Abdallah, F. Pattus, The Crystal Structure of the Pyoverdine Outer Membrane Receptor FpvA from *Pseudomonas aeruginosa* at 3.6 Å Resolution. J. Mol. Biol. 347, 121–134 (2005).

57. G. M. Singh, P. D. Fortin, A. Koglin, C. T. Walsh, β-Hydroxylation of the Aspartyl Residue in the Phytotoxin Syringomycin E: Characterization of Two Candidate Hydroxylases AspH and SyrP in *Pseudomonas syringae*. Biochemistry 47, 11310–11320 (2008).

58. J.-M. Meyer, Exogenous siderophore-mediated iron uptake in Pseudomonas aeruginosa: possible involvement of porin OprF in iron translocation. J. Gen. Microbiol. 138, 951–958 (1992).

59. E. Sugawara, E. M. Nestorovich, S. M. Bezrukov, H. Nikaido, Pseudomonas aeruginosa Porin OprF Exists in Two Different Conformations. J. Biol. Chem. 281, 16220–16229 (2006).

60. P. Nadal-Jimenez, G. Koch, C. R. Reis, R. Muntendam, H. Raj, C. M. Jeronimus-Stratingh, R. H. Cool, W. J. Quax, PvdP Is a Tyrosinase That Drives Maturation of the Pyoverdine Chromophore in Pseudomonas aeruginosa. J. Bacteriol. 196, 2681–2690 (2014).

61. M. T. Ringel, G. Dräger, T. Brüser, PvdO is required for the oxidation of dihydropyoverdine as the last step of fluorophore formation in *Pseudomonas fluorescens*. J. Biol. Chem. 293, 2330–2341 (2018).

62. B. J. McMorran, M. E. Merriman, I. T. Rombel, I. L. Lamont, Characterisation of the *pvdE* gene which is required for pyoverdine synthesis in *Pseudomonas aeruginosa*. Gene 176, 55–59 (1996).

63. E. J. Drake, A. M. Gulick, Structural Characterization and High-Throughput Screening of Inhibitors of PvdQ, an NTN Hydrolase Involved in Pyoverdine Synthesis. ACS Chem. Biol. 6, 1277–1286 (2011).

64. M. T. Ringel, G. Dräger, T. Brüser, PvdN Enzyme Catalyzes a Periplasmic Pyoverdine Modification. J. Biol. Chem. 291, 23929–23938 (2016).

65. M. Wehrmann, C. Berthelot, P. Billard, J. Klebensberger, Rare Earth Element (REE)-Dependent Growth of *Pseudomonas putida* KT2440 Relies on the ABC-Transporter PedA1A2BC and Is Influenced by Iron Availability. Front. Microbiol. 10 (2019).

66. T. Henríquez, N. V. Stein, H. Jung, PvdRT-OpmQ and MdtABC-OpmB efflux systems are involved in pyoverdine secretion in Pseudomonas putida KT2440. Environ. Microbiol. Rep. 11, 98–106 (2019).

67. N. V. Stein, M. Eder, F. Burr, S. Stoss, L. Holzner, H.-H. Kunz, H. Jung, The RND efflux system ParXY affects siderophore secretion in Pseudomonas putida KT2440. Microbiol. Spectr. 11, e02300–23 (2023).

68. N. V. Stein, M. Eder, S. Brameyer, S. Schwenkert, H. Jung, The ABC transporter family efflux pump PvdRT-OpmQ of Pseudomonas putida KT2440: purification and initial characterization. FEBS Lett. 597, 1403–1414 (2023).

69. M. Wehrmann, P. Billard, A. Martin-Meriadec, A. Zegeye, J. Klebensberger, Functional Role of Lanthanides in Enzymatic Activity and Transcriptional Regulation of Pyrroloquinoline Quinone-Dependent Alcohol Dehydrogenases in *Pseudomonas putida* KT2440. mBio 8, 10.1128/mbio.00570-17 (2017).

70. D. Kraemer, M. Bau, Siderophores and the formation of cerium anomalies in anoxic environments. Geochem. Perspect. Lett. 22, 50–55 (2022).

71. G. Tircsó, Z. Garda, F. K. Kálmán, Z. Baranyai, I. Pócsi, G. Balla, I. Tóth, Lanthanide(III) complexes of some natural siderophores: A thermodynamic, kinetic and relaxometric study. J. Inorg. Biochem. 127, 53–61 (2013).

72. I. J. Schalk, C. Hennard, C. Dugave, K. Poole, M. A. Abdallah, F. Pattus, Iron-free pyoverdin binds to its outer membrane receptor FpvA in Pseudomonas aeruginosa: a new mechanism for membrane iron transport. Mol. Microbiol. 39, 351–361 (2001).

73. H. Adams, G. Zeder-Lutz, I. Schalk, F. Pattus, H. Celia, Interaction of TonB with the Outer Membrane Receptor FpvA of *Pseudomonas aeruginosa*. J. Bacteriol. 188, 5752–5761 (2006).

74. P. Godoy, M. I. Ramos-González, J. L. Ramos, Involvement of the TonB System in Tolerance to Solvents and Drugs in *Pseudomonas putida* DOT-T1E. J. Bacteriol. 183, 5285–5292 (2001).

75. P. Godoy, M. Ramos-González, J. L. Ramos, *Pseudomonas putida* mutants in the *exbBexbDtonB* gene cluster are hypersensitive to environmental and chemical stressors. Environ. Microbiol. 6, 605–610 (2004).

76. P. Roszczenko-Jasińska, H. N. Vu, G. A. Subuyuj, R. V. Crisostomo, J. Cai, N. F. Lien, E. J. Clippard, E. M. Ayala, R. T. Ngo, F. Yarza, J. P. Wingett, C. Raghuraman, C. A. Hoeber, N. C. Martinez-Gomez, E. Skovran, Gene products and processes contributing to lanthanide homeostasis and methanol metabolism in Methylorubrum extorquens AM1. Sci. Rep. 10, 12663 (2020).

77. P. Cornelis, S. Matthijs, Diversity of siderophore-mediated iron uptake systems in fluorescent pseudomonads: not only pyoverdines. Environ. Microbiol. 4, 787–798 (2002).

78. J.-M. Meyer, V. A. Geoffroy, C. Baysse, P. Cornelis, I. Barelmann, K. Taraz, H. Budzikiewicz, Siderophore-mediated iron uptake in fluorescent Pseudomonas: characterization of the pyoverdine-receptor binding site of three cross-reacting pyoverdines. Arch. Biochem. Biophys. 397, 179–183 (2002).

79. C. E. Ugwu, O. C. Akinsulie, T. F. Ayandokun, F. A. Ajibade, S. Shahzad, V. A. Aliyu, M. J. Oladoye, I. Idris, K. O. Obasi, J. K. Edeh, A.-A. A. Olojede, C. B. Ukauwa, M. I. Adeyemi, C. C. Ugwu, L. C. Ugorji, Mechanisms and Therapeutic Potential of Nutritional Immunity. Pathogens 15, 176 (2026).

80. G. Ghssein, Z. Ezzeddine, A Review of Pseudomonas aeruginosa Metallophores: Pyoverdine, Pyochelin and Pseudopaline. Biology 11, 1711 (2022).

81. E. Butaitė, M. Baumgartner, S. Wyder, R. Kümmerli, Siderophore cheating and cheating resistance shape competition for iron in soil and freshwater Pseudomonas communities. Nat. Commun. 8, 414 (2017).

82. H. S. Zurier, C. Hart, S. Napieralski, W. R. Henson, S. Banta, K. H. Kucharzyk, Extremophilic microbes tolerate high concentrations of rare earth elements (REEs). Biotechnol. Lett. 47, 113 (2025).

83. Z. Ezzeddine, G. Ghssein, Metallophores as promising chelates for heavy metals removal from polluted water. Bioremediation J. 30, 379–381 (2024).

84. P. Rasoulnia, S. M. Mousavi, Maximization of organic acids production by *Aspergillus niger* in a bubble column bioreactor for V and Ni recovery enhancement from power plant residual ash in spent-medium bioleaching experiments. Bioresour. Technol. 216, 729–736 (2016).

85. M. Alipanah, H. Jin, Q. Zhou, C. Barboza, D. Gazzo, V. Thompson, Y. Fujita, J. Liu, A. Anderko, D. Reed, Sustainable bioleaching of lithium-ion batteries for critical metal recovery: Process optimization through design of experiments and thermodynamic modeling. Resour. Conserv. Recycl. 199, 107293 (2023).

86. N. M. Good, C. S. Kang-Yun, M. Z. Su, A. M. Zytnick, C. C. Barber, H. N. Vu, J. M. Grace, H. H. Nguyen, W. Zhang, E. Skovran, M. Fan, D. M. Park, N. C. Martinez-Gomez, Scalable and Consolidated Microbial Platform for Rare Earth Element Leaching and Recovery from Waste Sources. Environ. Sci. Technol. 58, 570–579 (2024).

87. A. M. Schmitz, B. Pian, S. Marecos, M. Wu, M. Holycross, E. Gazel, M. C. Reid, B. Barstow, High efficiency rare earth element bioleaching with systems biology guided engineering of Gluconobacter oxydans. *Commun*. Biol. 8, 815 (2025).

88. A. Jaiswal, S. I. Raj, A. Isiaka Adetunji, L. Negadi, S. Singh, K. Tumba, I. Bahadur, I. Uddin, Bioleaching as an Eco-Friendly Nano-Factory for Sustainable Inorganic Waste Management: Current Advancements, Challenges, and Opportunities. ChemistryOpen 14, e202500104 (2025).

89. G. J.-P. Deblonde, J. A. Mattocks, D. M. Park, D. W. Reed, J. A. Cotruvo, Y. Jiao, Selective and Efficient Biomacromolecular Extraction of Rare-Earth Elements using Lanmodulin. Inorg. Chem. 59, 11855–11867 (2020).

90. J. A. Mattocks, J. J. Jung, C.-Y. Lin, Z. Dong, N. H. Yennawar, E. R. Featherston, C. S. Kang-Yun, T. A. Hamilton, D. M. Park, A. K. Boal, J. A. Cotruvo, Enhanced rare-earth separation with a metal-sensitive lanmodulin dimer. Nature 618, 87–93 (2023).

91. C. W. Johnson, G. T. Beckham, Aromatic catabolic pathway selection for optimal production of pyruvate and lactate from lignin. Metab. Eng. 28, 240–247 (2015).

92. F. Meier, T. Williams, I. Paulsen, Welly: a web-tool for visualizing growth curves from microplate data. Bioinforma. Adv. 5 (2025).

93. R Core Team, R: A language and environment for statistical computing. (2021). https://www.r-project.org/.

94. H. Wickham, Ggplot2: Elegant Graphics for Data Analysis (Springer International Publishing, Cham, 2016; http://link.springer.com/10.1007/978-3-319-24277-4).

95. A. P. Arkin, R. W. Cottingham, C. S. Henry, N. L. Harris, R. L. Stevens, S. Maslov, P. Dehal, D. Ware, F. Perez, S. Canon, M. W. Sneddon, M. L. Henderson, W. J. Riehl, D. Murphy-Olson, S. Y. Chan, R. T. Kamimura, S. Kumari, M. M. Drake, T. S. Brettin, E. M. Glass, D. Chivian, D. Gunter, D. J. Weston, B. H. Allen, J. Baumohl, A. A. Best, B. Bowen, S. E. Brenner, C. C. Bun, J.-M. Chandonia, J.-M. Chia, R. Colasanti, N. Conrad, J. J. Davis, B. H. Davison, M. DeJongh, S. Devoid, E. Dietrich, I. Dubchak, J. N. Edirisinghe, G. Fang, J. P. Faria, P. M. Frybarger, W. Gerlach, M. Gerstein, A. Greiner, J. Gurtowski, H. L. Haun, F. He, R. Jain, M. P. Joachimiak, K. P. Keegan, S. Kondo, V. Kumar, M. L. Land, F. Meyer, M. Mills, P. S. Novichkov, T. Oh, G. J. Olsen, R. Olson, B. Parrello, S. Pasternak, E. Pearson, S. S. Poon, G. A. Price, S. Ramakrishnan, P. Ranjan, P. C. Ronald, M. C. Schatz, S. M. D. Seaver, M. Shukla, R. A. Sutormin, M. H. Syed, J. Thomason, N. L. Tintle, D. Wang, F. Xia, H. Yoo, S. Yoo, D. Yu, KBase: The United States Department of Energy Systems Biology Knowledgebase. Nat. Biotechnol. 36, 566– 569 (2018).

96. S. Andrews, FastQC: A Quality Control tool for High Throughput Sequence Data (2010). https://www.bioinformatics.babraham.ac.uk/projects/fastqc/.

97. A. M. Bolger, M. Lohse, B. Usadel, Trimmomatic: a flexible trimmer for Illumina sequence data. Bioinformatics 30, 2114–2120 (2014).

98. B. Langmead, S. L. Salzberg, Fast gapped-read alignment with Bowtie 2. Nat. Methods 9, 357–359 (2012).

99. M. Pertea, G. M. Pertea, C. M. Antonescu, T.-C. Chang, J. T. Mendell, S. L. Salzberg, StringTie enables improved reconstruction of a transcriptome from RNA-seq reads. Nat. Biotechnol. 33, 290–295 (2015).

100. M. I. Love, W. Huber, S. Anders, Moderated estimation of fold change and dispersion for RNA-seq data with DESeq2. Genome Biol. 15, 550 (2014).

101. M. C. Chambers, B. Maclean, R. Burke, D. Amodei, D. L. Ruderman, S. Neumann, L. Gatto, B. Fischer, B. Pratt, J. Egertson, K. Hoff, D. Kessner, N. Tasman, N. Shulman, B. Frewen, T. A. Baker, M.-Y. Brusniak, C. Paulse, D. Creasy, L. Flashner, K. Kani, C. Moulding, S. L. Seymour, L. M. Nuwaysir, B. Lefebvre, F. Kuhlmann, J. Roark, P. Rainer, S. Detlev, T. Hemenway, A. Huhmer, J. Langridge, B. Connolly, T. Chadick, K. Holly, J. Eckels, E. W. Deutsch, R. L. Moritz, J. E. Katz, D. B. Agus, M. MacCoss, D. L. Tabb, P. Mallick, A cross-platform toolkit for mass spectrometry and proteomics. Nat. Biotechnol. 30, 918–920 (2012).

102. S. Heuckeroth, T. Damiani, A. Smirnov, O. Mokshyna, C. Brungs, A. Korf, J. D. Smith, P. Stincone, N. Dreolin, L.-F. Nothias, T. Hyötyläinen, M. Orešič, U. Karst, P. C. Dorrestein, D. Petras, X. Du, J. J. J. Van Der Hooft, R. Schmid, T. Pluskal, Reproducible mass spectrometry data processing and compound annotation in MZmine 3. Nat. Protoc. 19, 2597–2641 (2024).

103. M. Wang, J. J. Carver, V. V. Phelan, L. M. Sanchez, N. Garg, Y. Peng, D. D. Nguyen, J. Watrous, C. A. Kapono, T. Luzzatto-Knaan, C. Porto, A. Bouslimani, A. V. Melnik, M. J. Meehan, W.-T. Liu, M. Crüsemann, P. D. Boudreau, E. Esquenazi, M. Sandoval-Calderón, R. D. Kersten, L. A. Pace, R. A. Quinn, K. R. Duncan, C.-C. Hsu, D. J. Floros, R. G. Gavilan, K. Kleigrewe, T. Northen, R. J. Dutton, D. Parrot, E. E. Carlson, B. Aigle, C. F. Michelsen, L. Jelsbak, C. Sohlenkamp, P. Pevzner, A. Edlund, J. McLean, J. Piel, B. T. Murphy, L. Gerwick, C.-C. Liaw, Y.-L. Yang, H.-U. Humpf, M. Maansson, R. A. Keyzers, A. C. Sims, A. R. Johnson, A. M. Sidebottom, B. E. Sedio, A. Klitgaard, C. B. Larson, C. A. Boya P, D. Torres-Mendoza, D. J. Gonzalez, D. B. Silva, L. M. Marques, D. P. Demarque, E. Pociute, E. C. O’Neill, E. Briand, E. J. N. Helfrich, E. A. Granatosky, E. Glukhov, F. Ryffel, H. Houson, H. Mohimani, J. J. Kharbush, Y. Zeng, J. A. Vorholt, K. L. Kurita, P. Charusanti, K. L. McPhail, K. F. Nielsen, L. Vuong, M. Elfeki, M. F. Traxler, N. Engene, N. Koyama, O. B. Vining, R. Baric, R. R. Silva, S. J. Mascuch, S. Tomasi, S. Jenkins, V. Macherla, T. Hoffman, V. Agarwal, P. G. Williams, J. Dai, R. Neupane, J. Gurr, A. M. C. Rodríguez, A. Lamsa, C. Zhang, K. Dorrestein, B. M. Duggan, J. Almaliti, P.-M. Allard, P. Phapale, L.-F. Nothias, T. Alexandrov, M. Litaudon, J.-L. Wolfender, J. E. Kyle, T. O. Metz, T. Peryea, D.-T. Nguyen, D. VanLeer, P. Shinn, A. Jadhav, R. Müller, K. M. Waters, W. Shi, X. Liu, L. Zhang, R. Knight, P. R. Jensen, B. Ø. Palsson, K. Pogliano, R. G. Linington, M. Gutiérrez, N. P. Lopes, W. H. Gerwick, B. S. Moore, P. C. Dorrestein, N. Bandeira, Sharing and community curation of mass spectrometry data with Global Natural Products Social Molecular Networking. Nat. Biotechnol. 34, 828–837 (2016).

104. A. K. Pakkir Shah, A. Walter, F. Ottosson, F. Russo, M. Navarro-Díaz, J. Boldt, J.-C. Kalinski, E. E. Kontou, J. Elofson, A. Polyzois, C. González-Marín, S. Farrell, M. R. Aggerbeck, T. Pruksatrakul, N. Chan, Y. Wang, M. Pöchhacker, C. Brungs, B. Cámara, A. M. Caraballo-Rodríguez, A. Cumsille, F. de Oliveira, K. Dührkop, Y. El Abiead, C. Geibel, L. G. Graves, M. Hansen, S. Heuckeroth, S. Knoblauch, A. Kostenko, M. CM. Kuijpers, K. Mildau, S. Papadopoulos Lambidis, P. W. Portal Gomes, T. Schramm, K. Steuer-Lodd, P. Stincone, S. Tayyab, G. A. Vitale, B. C. Wagner, S. Xing, M. T. Yazzie, S. Zuffa, M. de Kruijff, C. Beemelmanns, H. Link, C. Mayer, J. J. van der Hooft, T. Damiani, T. Pluskal, P. C. Dorrestein, J. Stanstrup, R. Schmid, M. Wang, A. T. Aron, M. Ernst, D. Petras, The Hitchhiker’s Guide to Statistical Analysis of Feature-based Molecular Networks from Non-Targeted Metabolomics Data. ChemRxiv, doi: 10.26434/chemrxiv-2023-wwbt0 (2023).

